# Deciphering the Molecular Mechanism of HCV Protease Inhibitor Fluorination as a General Approach to Avoid Drug Resistance

**DOI:** 10.1101/2021.11.30.470632

**Authors:** Jacqueto Zephyr, Desaboini Nageswara Rao, Sang V. Vo, Mina Henes, Klajdi Kosovrasti, Ashley N. Matthew, Adam K. Hedger, Jennifer Timm, Elise T. Chan, Akbar Ali, Nese Kurt Yilmaz, Celia A. Schiffer

## Abstract

Third generation Hepatitis C virus (HCV) NS3/4A protease inhibitors (PIs), glecaprevir and voxilaprevir, are highly effective across genotypes and against many resistant variants. Unlike earlier PIs, these compounds have fluorine substitutions on the P2-P4 macrocycle and P1 moieties. Fluorination has long been used in medicinal chemistry as a strategy to improve physicochemical properties and potency. However, the molecular basis by which fluorination improves potency and resistance profile of HCV NS3/4A PIs is not well understood. To systematically analyze the contribution of fluorine substitutions to inhibitor potency and resistance profile, we used a multi-disciplinary approach involving inhibitor design and synthesis, enzyme inhibition assays, co-crystallography, and structural analysis. A panel of inhibitors in matched pairs were designed with and without P4 cap fluorination, tested against WT protease and the D168A resistant variant, and a total of 22 high-resolution co-crystal structures were determined. While fluorination did not significantly improve potency against the WT protease, PIs with fluorinated P4 caps retained much better potency against the D168A protease variant. Detailed analysis of the co-crystal structures revealed that PIs with fluorinated P4 caps can sample alternate binding conformations that enable adapting to structural changes induced by the D168A substitution. Our results elucidate molecular mechanisms of fluorine-specific inhibitor interactions that can be leveraged in avoiding drug resistance.

## INTRODUCTION

Hepatitis C virus (HCV) infects about 71 million people worldwide and is responsible for 400,000 deaths per year (1). HCV infection eventually leads to chronic liver disease and is the leading cause of hepatocellular carcinoma (2). Current treatment involves a combination of direct-acting antivirals (DAAs) targeting viral proteins NS5A, NS5B and NS3/4A protease (3). The earlier generation HCV NS3/4A protease inhibitors (PIs) were readily susceptible to drug resistance and effective against only certain genotypes. These PIs were linear covalent peptidomimetics (telaprevir, boceprevir) or noncovalent P1-P3 macrocycles (simeprevir, paritaprevir). Currently, only three (grazoprevir, glecaprevir, and voxilaprevir) of the seven FDA-approved PIs are used in clinic (**Fig. 1a** and **S1**). All three of these PIs are P2-P4 macrocycles sharing a very similar chemical scaffold, with fluorine atoms incorporated at the P2^+^ and P1 moieties of glecaprevir (GLE) and voxilaprevir (VOX) (**Fig. 2**). These first pan-genotypic inhibitors are able to target the challenging genotype 3 (GT-3) NS3/4A protease, a milestone in HCV treatment.

**Figure 1.**
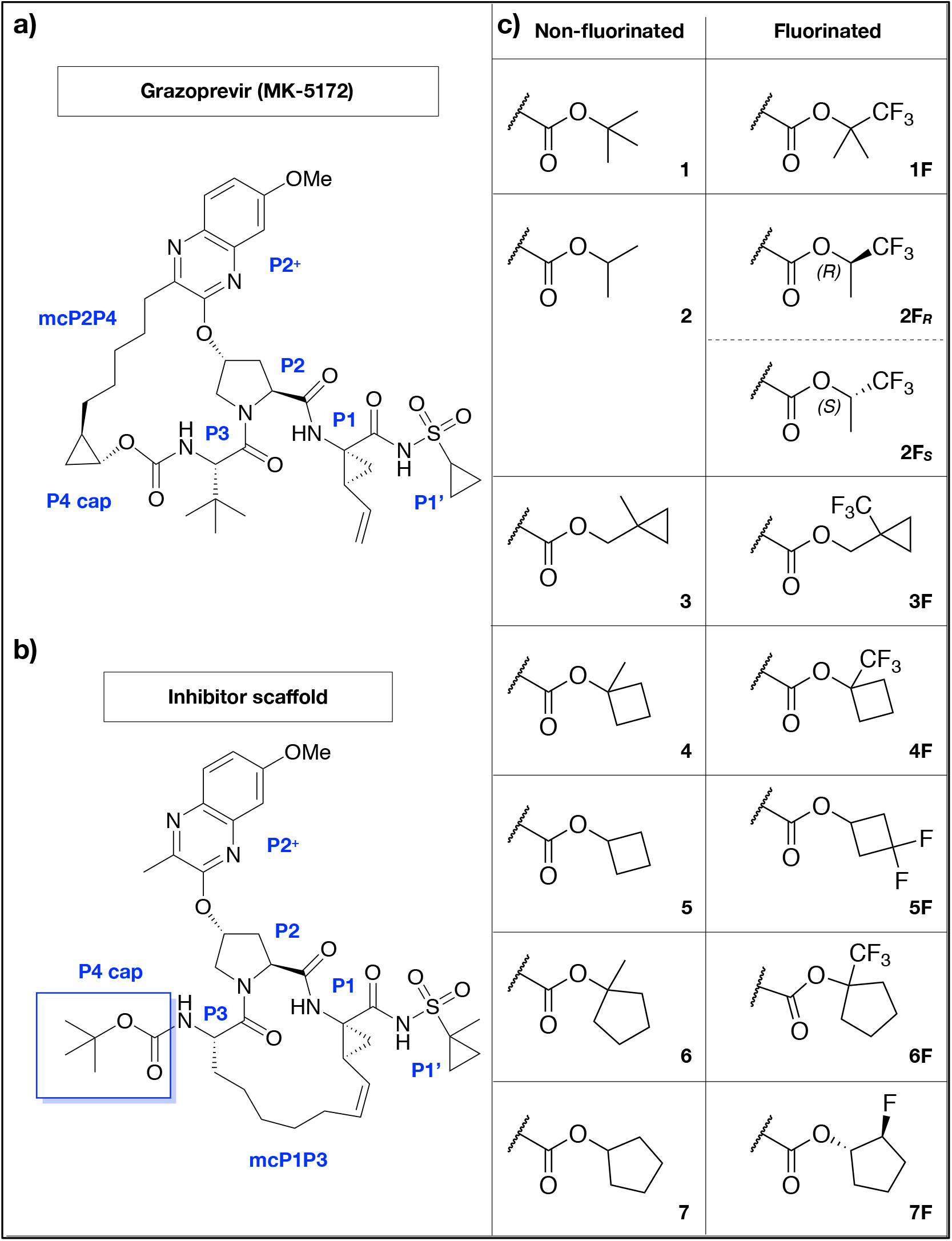
Inhibitor design strategy. **(a)** Grazoprevir, (**b**) P1-P3 macrocyclic inhibitor scaffold, and (**c**) non-fluorinated and fluorinated P4 capping groups used in this study. The moieties of grazoprevir and the inhibitor scaffold with macrocyclization between P2 and P4 (mcP2P4), and P1 and P3 (mcP1P3) side chains, respectively, are labeled.

**Figure 2.**
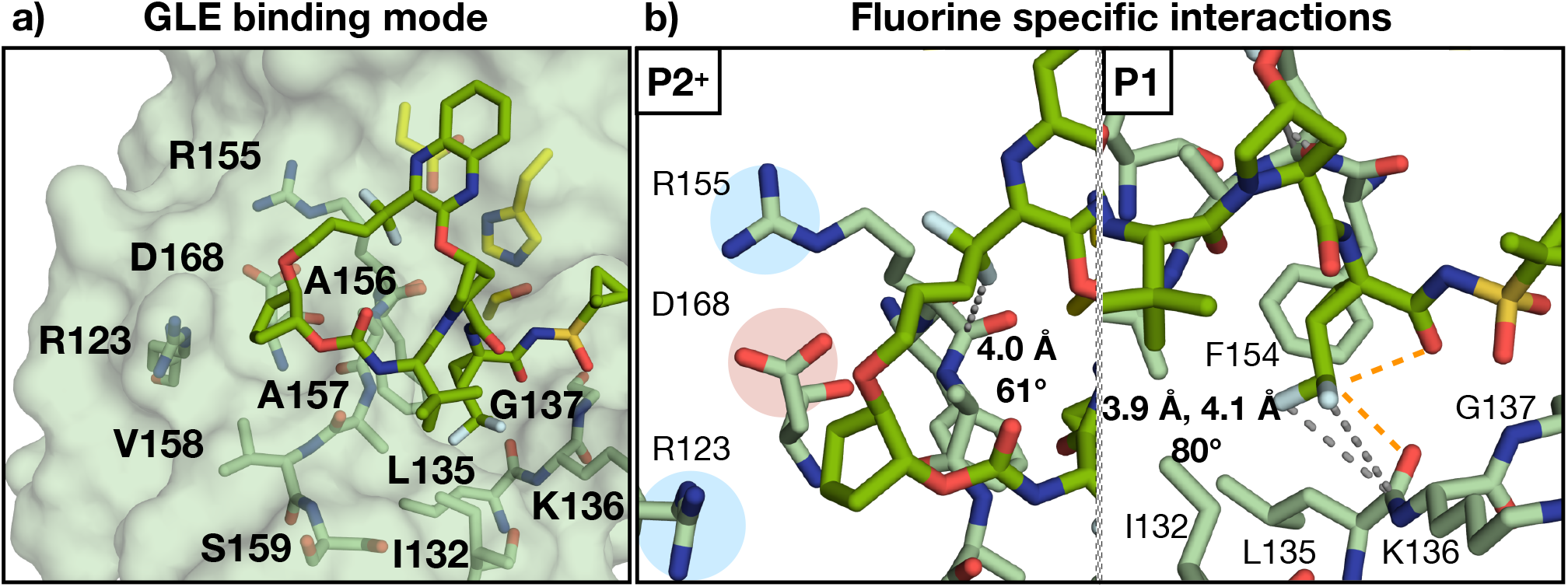
Structural analysis of Glecaprevir. **(a)** Overall binding mode of Glecaprevir to WT (PDB ID: 6P6L), and **(b)** fluorine specific interactions. Residues of the S4 pocket and around the fluorine atoms are labeled, gray dotted lines represent an orthogonal multipolar interaction (with distance and angle provided). The orange dotted lines (2.2 and 2.6 Å) represent fluorine induced hydrogen bonds of the P1 moiety. The electrostatic potential (blue for positive and red for negative) are highlighted.

Although GZR, GLE and VOX are pan-genotypic inhibitors, they are still susceptible to common resistance-associated substitutions (RASs), including A156T and D168A in GT1 protease (4). The P2-P4 macrocycles of GZR, GLE and VOX protrude beyond the HCV substrate envelope and clash with the larger Thr side chain in the A156T resistant variant (5). Loss of inhibitor potency is also caused by the D168Q polymorphism in GT-3 protease (6). GLE is robust against the D168Q polymorphism, but similar to GZR is highly susceptible to the D168A RAS which decreases the potency of all PIs by increasing protein dynamics and weakening inhibitor interactions in the S4 pocket (5, 7). The fluorine atoms in GLE and VOX are at least partially responsible for the improved potency across genotypes compared with GZR (8).

Incorporation of fluorine atoms in the scaffold of lead compounds can serve several purposes, such as improving potency, metabolic stability, physiochemical properties, and conformational selectivity (9-11). Furthermore, fluorination in drug discovery has been used as an approach to combat drug resistance in a variety of targets (12-17). The benefits of fluorination are associated with the strong electronegativity and relatively small atomic radius of fluorine. Hydrogen to fluorine substitution yields a carbon-fluorine bond that is highly polarized causing the fluorine atom to carry a partial negative charge (18). Recent studies have explored fluorination in HCV NS3/4A PIs GZR, asunaprevir, and simeprevir at the P1, P2^+^ and P4 moieties (8, 19-21). Overall, these studies show that fluorine incorporation improved potency and antiviral activity against resistant variants and across genotypes.

The HCV NS3/4A protease substrate envelope, which is defined as the consensus volume occupied by natural substrates, serves as a tool to understand the molecular basis of drug resistance and potency (22, 23). Relocating the macrocycle from P2-P4 to P1-P3 and staying within the substrate envelope we demonstrated that we could design inhibitors that avoid susceptibility to RASs at A156. Further modifications to the P1-P3 macrocyclic scaffold also ameliorated susceptibility to the D168A RAS by staying within the constraints of the protease substrate envelope and achieving shape complementarity with the contours of the S4 pocket (24).

Moreover, we recently published an extensive structure-activity relationship study (SAR) focusing on two series of compounds with different P2^+^ quinoxalines moiety in combination with diverse non-fluorinated and fluorinated P4 capping groups of varying size and shape (25). Our SAR results indicated that P4 capping groups that optimally fill the S4 pocket led to PIs with both excellent potencies and resistance profiles. Furthermore, incorporating fluorine motifs at the P4 capping groups was successful at improving potency against common resistant variants (D168A and A156T) and GT-3. Our strategy of using fluorinated P4 caps to target the variable S4 pocket proved effective, and emphasized the need for structural data to understand the contribution of fluorination to potency, resistance profile, and inhibitor binding mode.

In the current study, we investigate the impact of fluorination on molecular interactions underlying potency and resistance profile using a panel of NS3/4A PIs with and without fluorine substitutions through structural analysis of co-crystal structures (**Fig. 1b**). Seven sets of fluorinated and non-fluorinated analogues that differ by 1-3 hydrogen-to-fluorine substitution at the P4 capping groups were compared. The PIs share an identical P1-P3 macrocyclic scaffold containing a flexible quinoxaline moiety at the P2^+^ position and diverse acyclic and cyclic P4 capping groups(25). While all PIs lost potency due to the D168A RAS, PIs with fluorinated P4 cap groups retained better potency compared to the non-fluorinated analogues (**Fig. 3**). A total of 22 high-resolution co-crystal structures, 10 with the WT NS3/4A protease and 12 with the D168A variant, were determined and compared with our previously reported co-crystal structures of nonfluorinated analogues (24). Detailed structural analysis revealed that increased van der Waals (vdW) contacts, as well as electrostatic and fluorine-induced intramolecular interactions contribute to the improved potency of fluorinated PIs. When in complex with the D168A protease variant, PIs with fluorinated P4 caps sampled alternate binding conformations that enabled PIs to adapt to structural changes in the S4 pocket. The results provide insights into the molecular mechanism by which inhibitor fluorination can lead to improved robustness against drug resistance.

**Figure 3.**
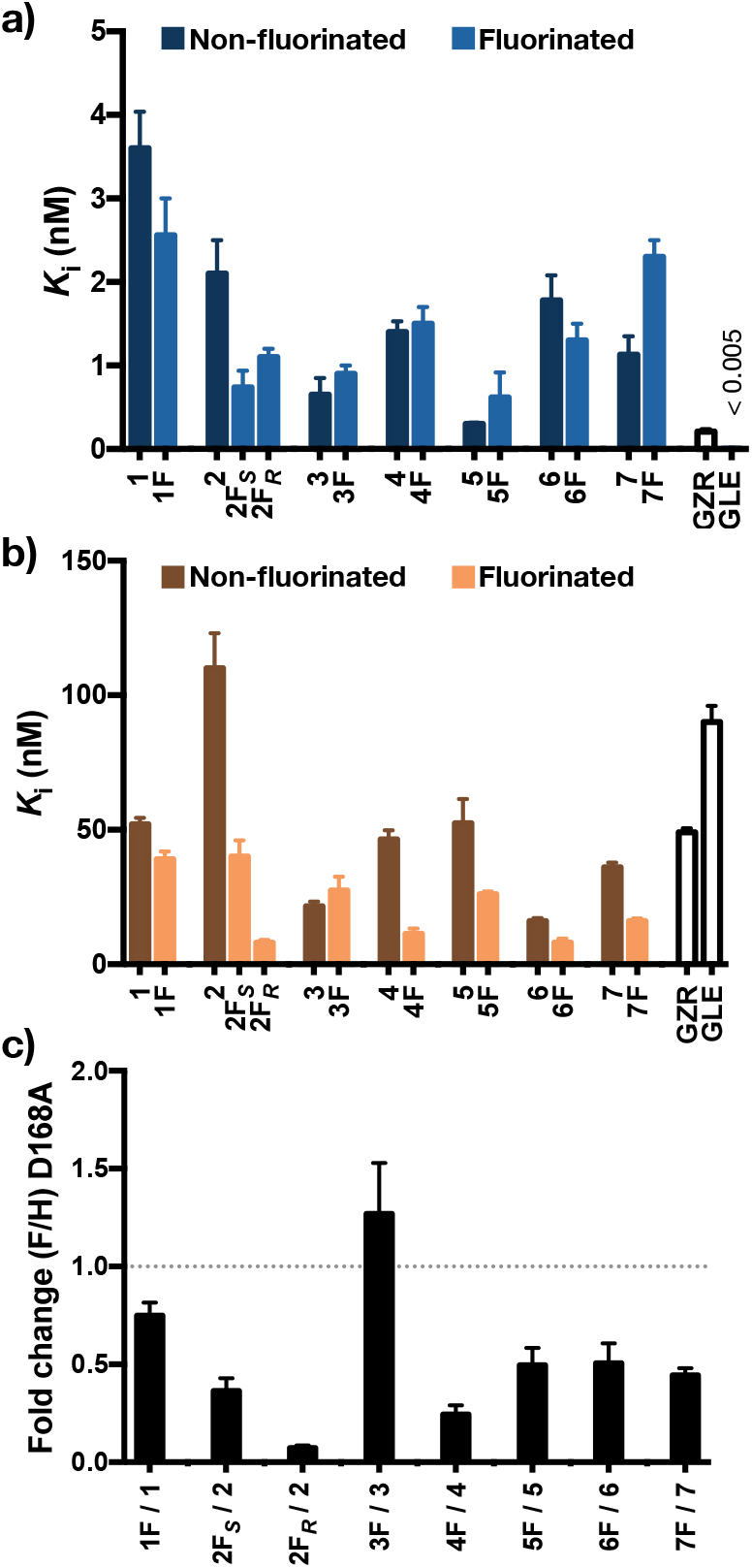
Inhibitor potencies and fold change analyses. Inhibition constants of PIs with non-fluorinated and fluorinated P4 caps, GRZ and GLE against **(a)** WT1a and **(b)** D168A variant. (**c**) Fold change in potency against the D168A resistant variant as a result of fluorination.

## RESULTS

### Fluorination improved potency against the D168A resistant variant

Seven sets of inhibitors were analyzed to determine the impact of fluorination on potency and efficacy against drug resistant variant D168A of HCV NS3/4A. The selected PIs contained non-fluorinated acyclic P4 capping groups *tert*-butyl (**1**), isopropyl (**2**), and cyclopropylethyl (**3**); and cyclic P4 capping groups 1-methylcyclocbutyl (**4**), cyclobutyl (**5**), 1-methylcyclopentyl (**6**), and cyclopentyl (**7**). The corresponding PIs with fluorinated acyclic P4 groups contain methyl to trifluoromethyl substitution, trifluoro tert-butyl (**1F**), trifluoro-isopropyl (**2F**_***S***_ and **2F**_***R***_,), and 1-methylcyclopropylethyl (**3F**) and with cyclic P4 groups contain either methyl to trifluoromethyl, 1-(trifluoromethyl)-cyclobutyl (**4F**) and 1-(trifluoromethyl)-cyclopentyl (**6F**) or hydrogen to fluorine substitutions, 3,3-difluorocyclobutyl (**5F**) and 2-fluorocyclopentyl (**7F**). These analogue pairs were selected because they differ only by 1-3 hydrogen to fluorine substitution and represent a set of chemically diverse P4 capping groups.

The PIs containing nonfluorinated P4 capping groups were potent against the WT protease with single digit nanomolar potencies (*K*_*i*_ = 0.3–3.6 nM) (**Fig. 3a**) (25). All these PIs lost significant potency against the D168A protease variant, ranging from 9- to 175-fold (**Fig. 3b, Fig. S2**, and **Table S1**). Overall, PIs with fluorinated P4 capping groups retained better potency relative to the respective non-fluorinated analogues (**Fig. 3b** and **c**). Fluorination of the cyclic P4 capping groups led to PIs with a 1.5- to 4.5-fold improvement in potency against the D168A protease variant (**Fig. S2a**). The distinct potency profiles of these PIs highlight the significance of fluorine substituent orientation to maintaining potency against drug resistant protease variants. As we have shown previously, PIs with bulky cyclic P4 caps that optimally fill and complement the contour of the S4 pocket maintain better potencies against the D168A protease variant (24). Addition of fluorine atoms to such P4 capping groups further increases inhibitor robustness. Overall, fluorine substitutions, especially in combination with larger cyclic P4 capping groups, were beneficial at retaining potency against the D168A protease variant.

To determine if the improvement in affinity observed for fluorinated inhibitors could be due to non-specific hydrophobic interactions, we assessed the lipophilicity of the compounds. As a measure of lipophilicity, the water:octanol distribution coefficients (LogD) were calculated (**Fig. S3**). The non-fluorinated inhibitors exhibited a correlation between lipophilicity and potency against the D168A protease variant (R^2^=0.66), in agreement with our previous observation that the inhibitors with larger P4 capping groups were more potent because of optimal packing in the S4 pocket (24). While overall the fluorinated inhibitors were more lipophilic (by 0.2-0.6 units) compared to their respective analogue, there was no correlation between LogD and potency against the WT protease or the D168A variant (**Fig. S3**). Therefore, fluorine-specific interactions are likely driving the improvement in potency.

### Crystal structures of protease–inhibitor complexes

To understand the contribution of fluorination to potency, co-crystal structures of both non-fluorinated and fluorinated PIs in complex with the WT NS3/4A protease and D168A variant were determined. A total of 22 new high-resolution co-crystal structures were solved, 10 in complex with the WT protease and 12 with the D168A protease variant, with resolutions ranging from 1.43 to 2.11 Å. The new structures were compared with previously determined structures of inhibitors **1, 6**, and **7** bound to the WT protease, and **6** and **4** bound to the D168A protease variant (PDB: 5VOJ, 6DIU, 6DIT, and 6PJ1and 6PIY respectively) (23, 24). Molecular models of **2** bound to the WT and D168A protease variant (see Methods section) were generated as diffraction quality crystals were not forthcoming. Altogether, these structures provide detailed comparison between the PIs containing non-fluorinated and fluorinated P4 capping groups. Specifically, the structures reveal that the fluorinated groups make fluorine-specific interactions with residues in the S4 pocket and either adopt an alternate conformation relative to the non-fluorinated P4 capping groups or sample two conformations.

### Fluorinated P4 caps can sample alternate conformations when bound to the D168A protease variant

The overall binding mode of PIs with fluorinated P4 caps were similar to those of the non-fluorinated inhibitors, with the P2^+^ quinoxaline moiety packing on the catalytic residues, as previously described (24). To gain insights into the ability of fluorines to cause major conformational changes in the binding mode of the P4 capping groups, co-crystal structures of each inhibitor pair were analyzed individually (**Fig. 1c**). More specifically, our analysis aimed at understanding the contribution of fluorine substitution to improved potency independent of the P4 capping group size (24).

Analysis of the first inhibitor pair revealed a distinct binding conformation of the fluorinated analogue only when bound to the D168A protease variant. **1F** binds to the WT protease in an overall similar binding conformation as **1**, with the *tert*-butyl and trifluoro *tert*-butyl P4 cap groups superimposing extremely well, and orienting toward R123 (**Fig. s4a, c**, and **g**). However, in the D168A protease variant, the trifluoro *tert*-butyl P4 cap group of **1F** samples two conformations. One conformation still orients the P4 capping group toward R123, while in the alternate conformation the entire P4 capping group is rotated and the trifluoromethyl moiety points toward R155 (**Fig. s4b, d, e**, and **h**). In this alternate conformation, the fluorinated P4 cap occupies the space created by the D168A mutation in the S4 pocket, likely contributing to the better potency compared to **1**, as filling the S4 pocket can lead to improved potency (**Fig. s4f**) (24). The absence of such an alternate binding conformation of **1F** when bound to the WT protease is likely due to repulsive interactions between the fluorine atoms and the carboxylic side chain of D168 and not due to a lack of space in the S4 pocket as bulkier P4 capping groups can be accommodated.

To further understand the contribution of the alternate conformation observed in the co-crystal structure of **1F** bound to the D168A protease variant, we determined co-crystal structures of the trifluoro isopropyl P4 capping group containing inhibitors, **2F**_***S***_ and **2F**_***R***_, that were designed to orient the fluorine atoms either toward the protein surface or solvent (**Fig. 1**). The co-crystal structures revealed that the trifluoromethyl group in the *S*-configuration was oriented toward solvent, while the one with the *R*-configuration was oriented toward the S4 pocket (**Fig. s5**). In the complex structures with WT and D168A protease variant, the P4 cap methyl group of **2F**_***S***_ pointed directly at residue 168 while the trifluoromethyl group was solvent exposed. Although the conformations of **2F**_***S***_ and **2F**_***R***_ bound to the WT and D168A protease variant were near identical, the side chain of R123 adopted distinct conformations. In the WT protease, the guanidinium side chain of R123 participated in an extensive electrostatic network with D168 and R155, and packed against the side chain of V158, which forms part of the S4 pocket. Meanwhile, in the complex structure of the D168A resistant variant, R123 adopted an alternate rotamer leading to a loss of vdW interactions with V158 which reshaped the S4 pocket. Despite having the P4 cap bound in opposite orientations, **2F**_***S***_ and **2F**_***R***_ were equipotent against the WT, but not the D168A protease. **2F**_***R***_, which had the trifluoromethyl moiety oriented toward the alanine at position 168, was 5-fold more potent compared to **2F**_***S***_ where the trifluoromethyl was solvent exposed. Notably, both **2F**_***S***_ and **2F**_***R***_ were significantly more potent against the D168A protease variant compared to the non-fluorinated analogue **2**, indicating that the fluorinated analogues had more favorable interactions with the D168A protease compared to the WT protease. This suggests that the carboxylic acid side chain of D168 in the WT protease may prevent the fluorine atoms from contributing maximally to potency. The significant improvement in potency against the D168A protease variant as a result of methyl to trifluoromethyl substitution demonstrates the potential of fluorination as a strategy to target resistant variants while maintaining potency toward the WT protease.

Unlike inhibitor pairs **1** and **1F, 3** and **3F** bind to the WT or D168A protease variant in a near identical binding mode. The 1-methyl substituent of the 1-(methyl) cyclopropylethyl P4 cap of **3** pointed toward D168 and the cyclopropyl moiety was solvent exposed (**Fig. s6**). The P2 moiety shifted toward the catalytic histidine in the complex structure with the D168A protease variant compared to that with the WT protease, with additional conformational changes in R123, R155, V158 and S159. Specifically, the distance between R123 and V158 increased by 2 Å and R155 sampled two conformations. Widening of the S4 pocket due to the D168A RAS allows the 1-methylcyclopropylethyl P4 cap of **3** to pack slightly deeper into to S4 pocket. Inhibitors **3** and **3F** were equipotent against both WT and D168A protease variant, indicating that 1-trifluoromethyl substituent at the P4 cap of **3F** likely does not contribute to binding affinity. This inhibitor pair represents the only case where fluorination failed to improve potency against the D168A protease variant.

### P4 cap fluorination can induce conformational changes distal from the S4 pocket

We next investigated the structural changes associated with fluorination of cyclic P4 cap groups containing PIs **4** and **5**. Inhibitor **4** was 4-fold less potent against the WT protease compared to **5**, despite having a larger P4 capping group. However, both inhibitors had similar potencies against the D168A resistant variant. The 1-methylcyclobutyl and cyclobutyl P4 capping groups of **4** and **5** respectively, bound to the WT and D168A protease variant with similar poses. The fluorinated analogue **4F**, with a 1-(trifluoromethyl)-cyclobutyl P4 cap bound in a conformation distinct from **4** in the co-crystal structures with the D168A protease variant (**Fig. s7**). The 1-trifluoromethyl substituent of **4F** was oriented toward R123 while the cyclobutyl moiety oriented toward the P2^+^ quinoxaline. Thus, both the 1-trifluoromethyl and the cyclobutyl moieties of the P4 cap interact with the protease, which differs from the binding mode of **4** where the 1-methyl substituent of the P4 cap is solvent exposed. Compared to the binding mode of the P4 cap of **4**, that of **4F** was rotated about the carbamate bond. The crystal structure of **4F** displayed additional differences beyond the P4 capping group, including a 1.2 Å shift of the P2^+^ quinoxaline moiety toward the catalytic histidine to avoid intramolecular steric clash with the P4 capping group. Protease residues around the S2^+^ subsite also experienced structural changes. V78, D79 and R155 sampled an alternate rotamer that reshaped the S2^+^ subsite while Y56, a clinically relevant RAS site, shifted toward the catalytic histidine (**Fig. s7**) (26). These structural changes allow the protease to maintain interactions with the quinoxaline moiety of **4F** that has been shifted relative to that of **4** (**Fig. s8**). These structural changes likely underlie the 4-fold improvement in potency of **4F** against the D168A protease variant compared to **4**. Therefore, fluorination allowed inhibitors to retain potency by enhancing contacts with the protease and causing distal structural changes from afar.

Optimally filling the S4 pocket of the D168A protease variant is critical for inhibitor potency. A 0.7 Å shift of the P4 capping group of **5** out of the S4 pocket in the complex structure with the resistant variant likely contributes to the 180-fold loss in potency compared to WT (**Fig. s9**) (24). In the complex structures with WT and D168A protease variant, one of the germinal fluorine atoms of the 3,3-difluorocyclobutyl P4 cap oriented toward the S4 pocket while the other was solvent exposed. However, the P4 cap cyclobutyl ring adopted a distinct ring pucker when bound to the D168A protease variant, which rotated the fluorine substituents ∼45° further into the S4 pocket and positioned one of the fluorine atoms at the center of the S4 pocket. The non-fluorinated P4 cap of **5** was elevated out of the S4 pocket in the D168A protease variant, while the fluorinated P4 cap of **5F** was oriented to fill the space created by the Asp to Ala mutation. This variation in the binding mode of non-fluorinated and fluorinated P4 caps of **5** and **5F**, respectively, may explain the 2-fold difference in potency between these two PIs against the D168A protease variant. The comparison of these complex structures shows that the non-fluorinated P4 capping groups were elevated out of the S4 pocket in the D168A protease resulting in loss in potency. Thus fluorine incorporation can drive P4 cap groups to reorient and improve contacts in the S4 pocket thereby helping PIs to retain potency.

As described in our previous studies, the cyclopentyl ring pucker complements the shape of the S4 pocket and makes extensive contacts with R123, A156, V158, and D/A168 (24). The binding conformation of **6** to the WT and D168A protease variant were identical, with the 1-methyl substituent at the P4 cap solvent exposed and the cyclopentyl moiety occupying the S4 pocket (**Fig. 4**). **6F** with a 1-(trifluoromethyl)-cyclopentyl P4 cap binds to the WT and D168A protease variants in two conformations where the lowest occupied conformation is similar to that of **6**. While in the alternate, higher occupancy conformation, the 1-trifluoromethyl substituent orients towards the guanidinium group of R123 with the cyclopentyl moiety positioned toward the P2^+^ quinoxaline. The alternate binding mode of the P4 cap of **6F** allows both the trifluoromethyl and cyclopentyl moieties to interact with the protease, and mirrors the binding mode of **4F** bound to the D168A protease variant.

**Figure 4.**
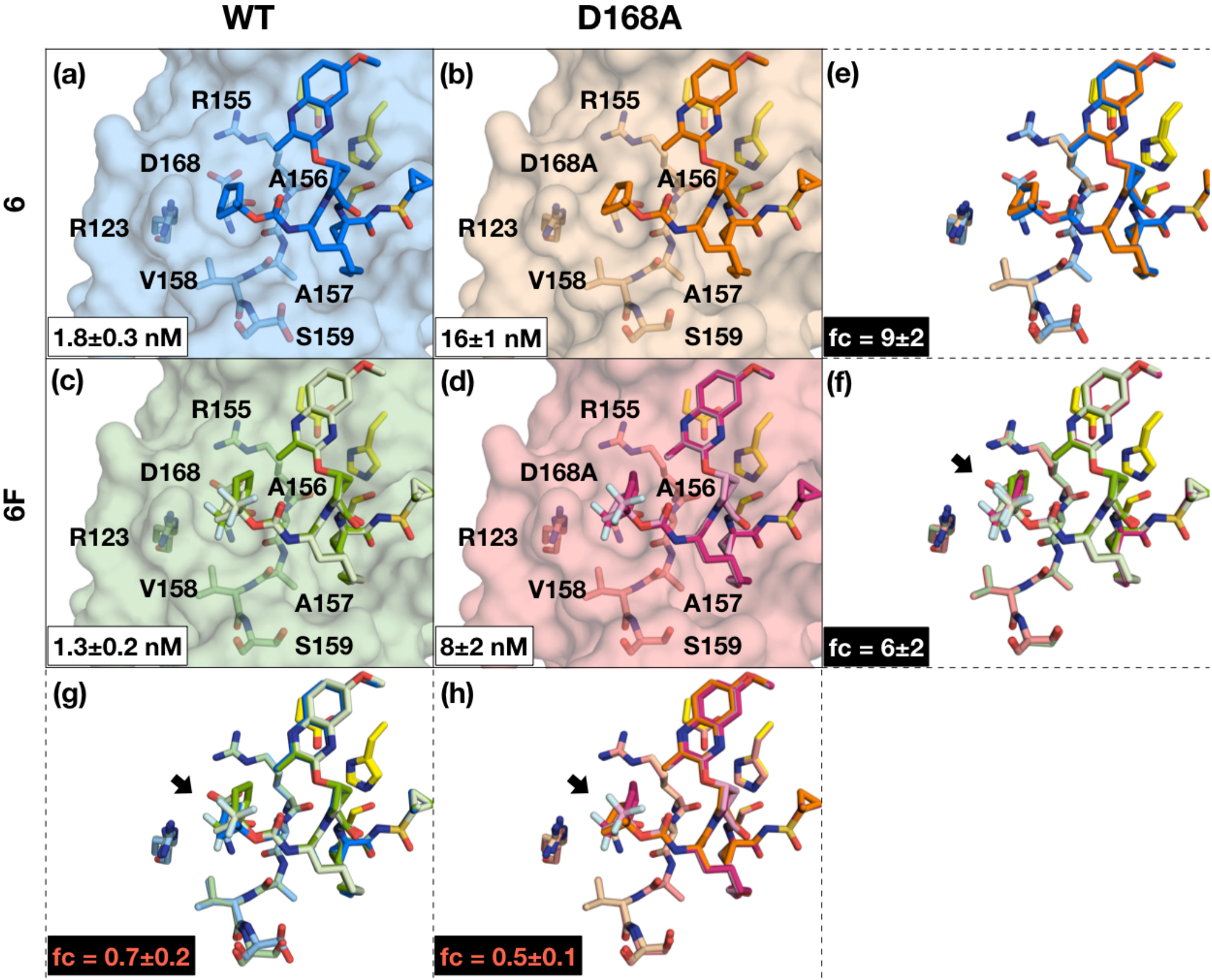
Co-crystal structures of **6** bound to WT1a **(a)** and D168A **(b)**, and **6F** bound to WT **(c)** and D168A **(d)**. The binding affinity is indicated in the respective panels **(a-d)**. Structural differences in **6** and **6F** binding to WT and D168A are shown in **(e)** and **(f)** respectively. Structural differences between **6** and **6F** binding to WT and D168A is shown in **(g)** and **(h)** respectively. The fold change (fc) in binding affinity relative to WT is indicated in the respective panels **(e** and **f)**. The fold change in binding affinity relative to the parent inhibitor is indicated in **(g)** and **(h)** for WT for D168A, respectively. Empty panels indicate no structural differences. The inhibitors and residues of the catalytic triad and S4 pocket (labeled) are shown as sticks. Inhibitors with two conformations are shown in a darker (conformation 1) and lighter (conformation 2) shade, where the darker shade represent the highest occupancy conformation. Arrows in panels **e**-**h** indicate structural differences in between the superimposed structures.

### Repulsive interactions of fluorinated P4 capping groups

As the cyclopentyl P4 cap proved to be optimal for targeting the S4 pocket, we investigated **7F** with a 2-fluorocyclopentyl P4 cap as the fluorinated analogue of **7** (**Fig. s10**). Binding of **7F** to both protease variants resulted in a 2 Å widening of the S4 pocket by increasing the distance between the side chains of R123 and V158, similar to the complex structure of **3** bound to the D168A protease variant. The 2-fluorocyclopentyl cap of **7F** bound in two conformation in WT protease complex structure, but adopted a single conformation in the D168A protease. In the highest occupied conformation in the WT protease, the fluorine atom is situated directly above the β carbon of D168. Notably, in this conformation, the 2-fluorocyclopentyl cap is elevated out of the S4 pocket by 0.9 Å relative to the binding conformation of the cyclopentyl cap of **7**, and rotated about the carbamate bond toward position 168. In the alternate and lower occupancy conformation, the 2-fluorocyclopentyl cap adopts a binding mode similar to the P4 cap of **7** with the fluorine atom solvent exposed, despite being highly unfavorable (27). This alternate conformation may explain the 2-fold loss in potency against the WT protease compared to the non-fluorinated analogue **7**. In the D168A protease structure, the 2-fluorocyclopentyl of **7F** adopts a single conformation that is identical to the higher occupancy conformation in the structure with the WT protease. The single conformation adopted by **7F** may explain 2-fold improvement in potency against the D168 variant compared the non-fluorinated analogue **7**. Overall, the distinct differences between the binding mode of **7F** in complex with WT and D168A protease variant indicate a repulsive interaction between the carboxylic sidechain of Asp at position 168 in the WT protease and the fluorine substituents of **7F**.

Generally across the respective pairs of non-fluorinated and fluorinated PIs, fluorination did not significantly improve potency against the WT protease but led to a ∼2-fold improvement against the D168A variant. Such observations, in combination with the binding mode of the fluorinated P4 caps suggest a repulsive interaction between fluorine and the carboxylic side chain of D168. The D168A RAS, however, eliminates such a repulsive interaction, allowing the fluorine atoms to optimally fill the space in the S4 pocket and establish improved contacts with the protease.

Overall, these sets of complex structures illustrate the impact of fluorination on the binding mode of the P4 capping group. Fluorinated PIs, **1F, 4F, 5F**, and **6F**, that sample alternate binding conformations in structures with the D168A protease variant are associated with a ∼2-fold better potency against this variant compared to the respective non-fluorinated analogue. Structural changes in the protease due to P4 capping group fluorination mainly occurred at residues R123, R155, and V158 around the S4 pocket. Additionally, we observed that the fluorinated PIs **1F, 6F**, and **7F** adopted two distinct conformations in the crystal structures, one that is similar to the conformation of the respective non-fluorinated analogue while the other differing by rotation of the P4 capping group. The freely rotatable carbamate bond linking the P4 cap to the rest of the inhibitor scaffold allows fluorinated P4 cap groups to rotate and sample an alternate conformation in the co-crystal structures. The non-fluorinated P4 cap groups also have the same conformational freedom but do not sample alternate binding conformations, indicating the critical role of fluorine in influencing the binding orientation of P4 caps.

### Alternate conformations of fluorinated inhibitors retain more vdW contacts

To understand the molecular interactions contributing to the improved potency of PIs with fluorinated P4 caps against the D168A protease variant, we calculated the vdW contact energies for each protease-inhibitor complex (**Fig. 5a** and **b**). Interestingly, the total vdW contact energies from WT protease and D168A variant structures follow the same trend as the *K*_i_ measurements, with **4F** and **7F** being the exceptions. Overall, the vdW analysis is in agreement with prior work showing that optimized packing in the S4 pocket of the D168A protease variant is required to maintain potency (24). As expected, all PIs lost vdW contacts due to the D168A mutation, consistent with our previous findings (23) (**Fig. 5c**). However, the alternate conformation sampled by the fluorinated PIs evaded the loss in vdW contacts and retained a vdW profile similar to when bound to the WT protease (**Fig. 5c**). These observations indicate that vdW contacts within the S4 pocket is a major contributor to retaining potency against D168A protease variant.

**Figure 5.**
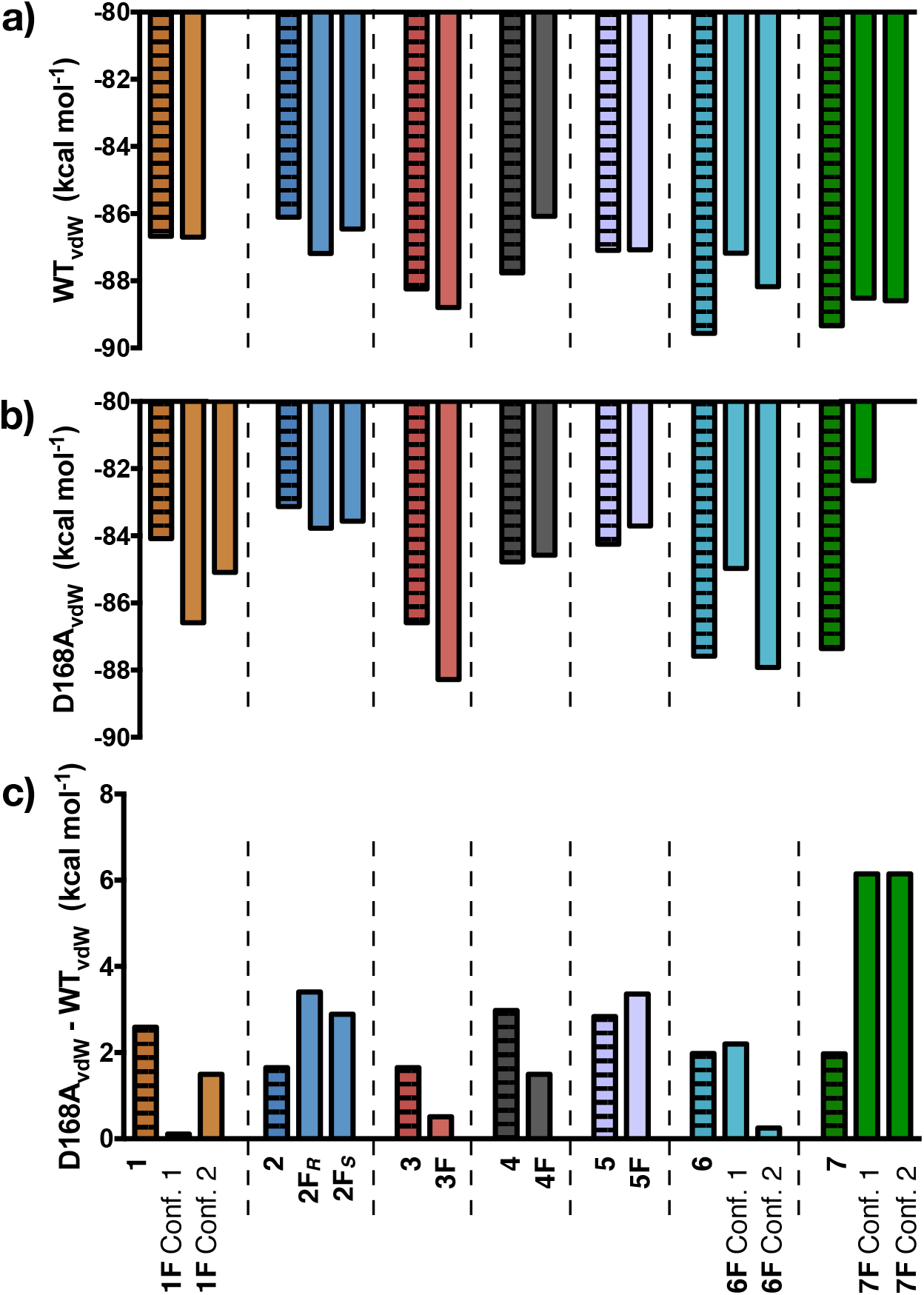
Loss of van de Waals contacts due to the D168A RAS. The total vdW contacts of the inhibitors bound to (**a**) WT and (**b**) D168A, and (**c**) the differences in vdW contacts between WT1a and D168A variant. The stripped pattern indicates parent inhibitor data.

Per residue vdW contact energies were calculated to identify the residues in the S4 pocket that lost contacts due to the D168A RAS. The loss of vdW contacts primarily and consistently occurred at position 168 and 156 for all inhibitors (**Fig. s11**). Notably, **2F**_***S***_ lost more vdW contacts compared to **2F**_***R***_, concurring with the inhibition data (**Fig. s11b**). The missing side chain for residue R123 in some of the structures causes discrepancies in vdW contacts for this specific residue (**Fig. s11e**). The large loss in vdW contact energies seen for **7F** is due to the Asp to Ala mutation and the widened S4 pocket (**Fig. s11g**). In summary, loss in vdW contacts at positions 156 and 168 are partly responsible for the observed loss in potency against the D168A protease variant.

To understand the contribution of fluorination to the vdW contact energies, we analyzed the differences in total vdW contacts of non-fluorinated and fluorinated PI pairs. When bound to the WT protease, fluorinated PIs did not lead to improved vdW contacts compared to non-fluorinated analogues, except for **2F**_***R***_ (**Fig. S12a**). However, in the complex structures with the D168A protease variant, **1, 2**, and **3** significantly benefited from fluorine substitutions (**Fig. S12b**). Overall, PIs with fluorinated P4 cap that adopt a single binding conformation had vdW contacts similar to that of the respective non-fluorinated analogue. Thus, the alternate binding conformations sampled by the PIs with fluorinated P4 caps had improved vdW contacts.

### Fluorine-specific interactions underlie inhibitor potency and binding conformation

To further understand the contribution of fluorine substitutions to inhibitor potency and binding mode, we analyzed fluorine-specific interactions. Fluorine atoms of all the fluorinated P4 caps formed favorable electrostatic interactions with the guanidinium side chain of R123, and appeared to have repulsive interaction with the carboxylic side chain of D168 in the WT protease (**Fig. 6**). The fluorine atoms also formed favorable orthogonal multipolar interactions with the imine carbon of R123 (**Fig. 6a**) (28-30). Orthogonal multipolar interaction between fluorine and amide carbonyl is estimated to contribute ∼0.2–0.4 kcal mol^-1^ to binding, while here the interaction is with the imine moiety, the calculated change in Gibbs binding free energy (Δ ΔG) from the measured *K*_*i*_ data against the D168A protease variant agrees with that estimation. Conversely, when targeting the WT protease, there is no clear contribution to binding energies from P4 cap fluorination (**Fig. S13**). Therefore, orthogonal multipolar interactions contribute to the potency of fluorinated PIs only against the D168A protease variant.

**Figure 6.**
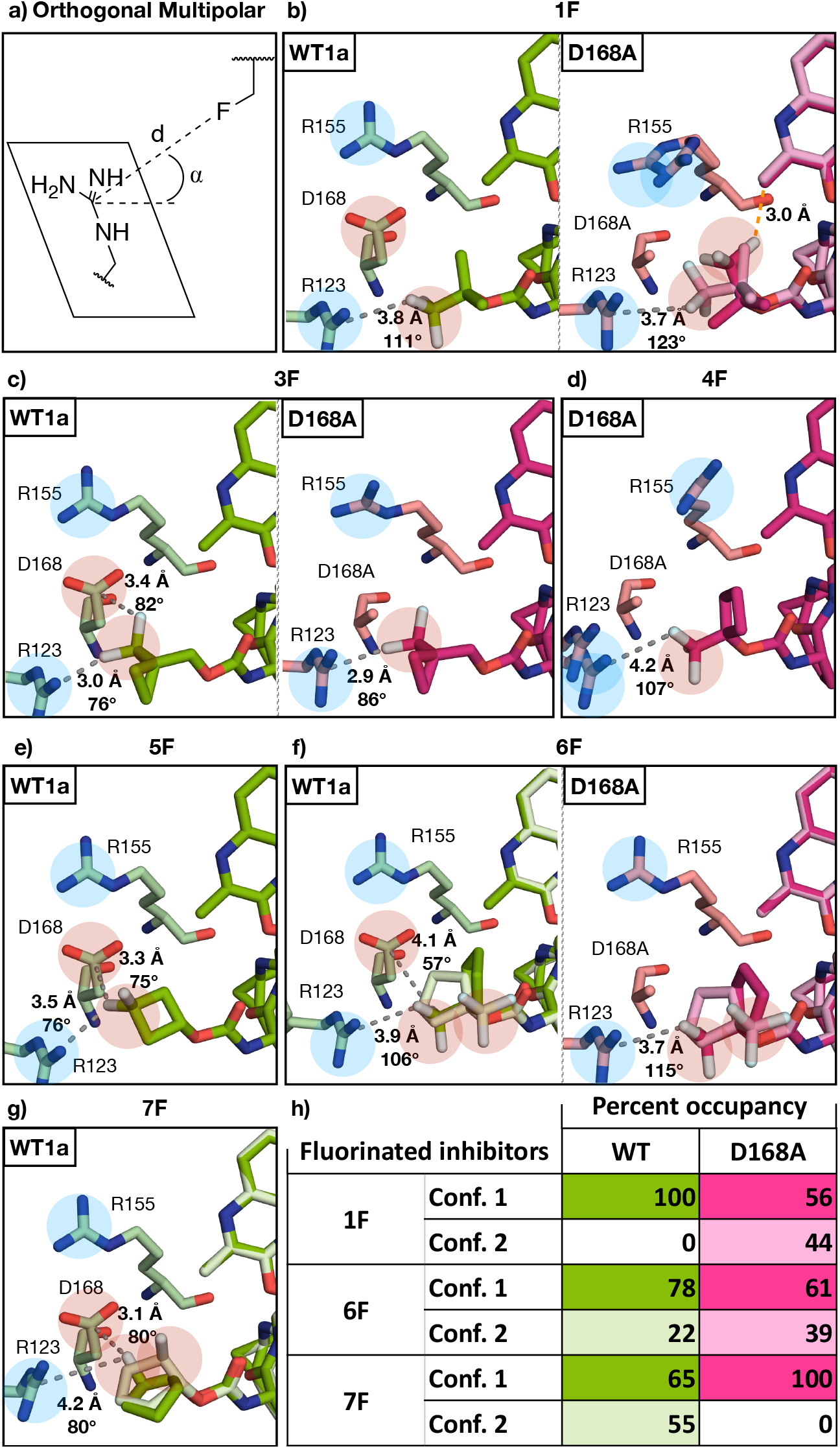
Fluorine specific interactions. **(a)** Geometric parameters of the orthogonal multipolar interaction as observed in the co-crystal structures with **(b) 1F, (c) 3F, (d) 4F, (e) 5F, (f) 6F**, and **(g) 7F. (h)** Percent occupancy for inhibitors sampling two conformations. Residues around the fluorinated moiety are labeled, gray dotted lines represent an orthogonal multipolar interaction (with distance and angle provided). The orange dotted line represents intramolecular fluorine hydrogen bonding with the acidic benzylic hydrogens. The electrostatic potential (blue for positive and red for negative) of the side chains and fluorine atoms are highlighted.

Furthermore, the high electronegativity of fluorine makes it possible to act as an hydrogen bond acceptor to acidic hydrogens (31). In the co-crystal structure of **1F** bound to the D168A protease variant, we observed an intramolecular fluorine hydrogen bonding interaction between one of the P4 cap trifluoro *tert*-butyl fluorine atoms and one of the acidic benzylic hydrogens (**Fig. 6b**). This intramolecular fluorine hydrogen bond interaction may be partly responsible for the two conformations observed in the co-crystal structures of **1F**. However, in the WT protease, the D168 side chain hinders the trifluoro *tert*-butyl P4 cap group of **1F** from adopting a conformation suitable for such an intramolecular interaction, indicating that fluorinated P4 caps are not maximizing all the possible fluorine-specific interactions when bound to the WT protease. Alternatively, D168 may influence PIs with fluorinated P4 capping groups to sample two conformations, as seen in co-crystal structure of **7F** bound to the WT protease, where the lower occupancy conformation is most probably sampled to mitigate the repulsive interaction between fluorine and the carboxylic side chain (**Fig. s10c** and **5g**).

In addition to affecting the conformations sampled by the fluorinated P4 caps, the identity of the residue at 168 can also affect the percent occupancies of inhibitors bound in multiple conformations. Although the two conformations of **6F** bound to the WT were identical to those in the D168A variant, the occupancy of these conformations varied (**Fig. 6f** and **h**). The occupancy of the alternate binding conformation increased by ∼2-fold (or ∼20%) in the D168A variant. Thus, the P4 cap of **6F** sampled the alternate conformation twice as frequently in the D168A variant, suggesting that the interaction with Asp at position 168 does not favor that alternate binding conformation. Hence, fluorine atoms of the P4 capping groups can act as sensors of the Asp to Ala RAS in the S4 pocket.

Fluorinated PIs bound to the WT protease prefer a conformation that avoids the repulsive interaction between fluorine and the D168 carboxylic side chain. In the D168A protease variant, the absence of such a repulsive interaction allows PIs with fluorinated P4 caps to adapt to the electrostatic and structural changes in the S4 pocket by sampling an alternate, more favorable conformation with improved vdW contacts.

Overall, the detailed structural analysis of 27 co-crystal structures and 3 modeled structures helped to elucidate the molecular mechanism by which PIs with fluorinated P4 caps retain potency against the D168A protease variant. Additionally, the intramolecular interaction observed between the P4 cap fluorine and the benzylic hydrogen of the P2^+^ quinoxaline, in the co-crystal structure of **1F** bound to the D168A protease variant, provides a strategy to further stabilize the P1-P3 macrocyclic scaffold toward improving potency against all genotypes and known resistant variants. In summary, the structural differences between PIs with and without and fluorinated P4 caps are driven by an ensemble of molecular interactions; (i) increase in vdW contacts, (ii) formation of orthogonal multipolar interaction with R123, (iii) avoidance of electrostatic repulsive with D168, and (iv) formation of intramolecular interactions (**Fig. 5** and **6**).

### Interdependence between P2^+^ and P4 capping groups

Two PIs, **8F** and **9F**, from our recent extensive SAR focusing on the P2^+^ quinoxaline moiety in combination with diverse non-fluorinated and fluorinated P4 capping groups, provide another strategy for designing intramolecular interaction between P2^+^ and P4 moieties (**Fig. 7**) (25). Although **8F** and **9F** contain a 6-methoxy-3-(trifluoromethyl)-quinoxaline P2^+^ moiety, which differs from the 7-methoxy-3-(methyl)-quinoxaline P2^+^ moiety, the P4 capping groups are identical to that of **1F** and **4F**, respectively. Inhibitor **8F** binds to the D168A protease variant in a conformation similar to that of **1F** with the P2^+^ quinoxaline moiety stacking on the catalytic triad. However, unlike **1F**, the trifluoro *tert*-butyl P4 capping group of **8F** adopts a single conformation with the trifluoromethyl moiety oriented towards R123. Potential dipole-dipole repulsion with the trifluoromethyl on the quinoxaline may hinder the trifluoro *tert*-butyl P4 capping group from sampling an alternate conformation similar to that observed in the complex structure of **1F** bound to the D168A protease variant (18). The trifluoromethyl substituent of the P2^+^ quinoxaline moiety therefore can influence the binding conformation of the fluorinated P4 capping group.

**Figure 7.**
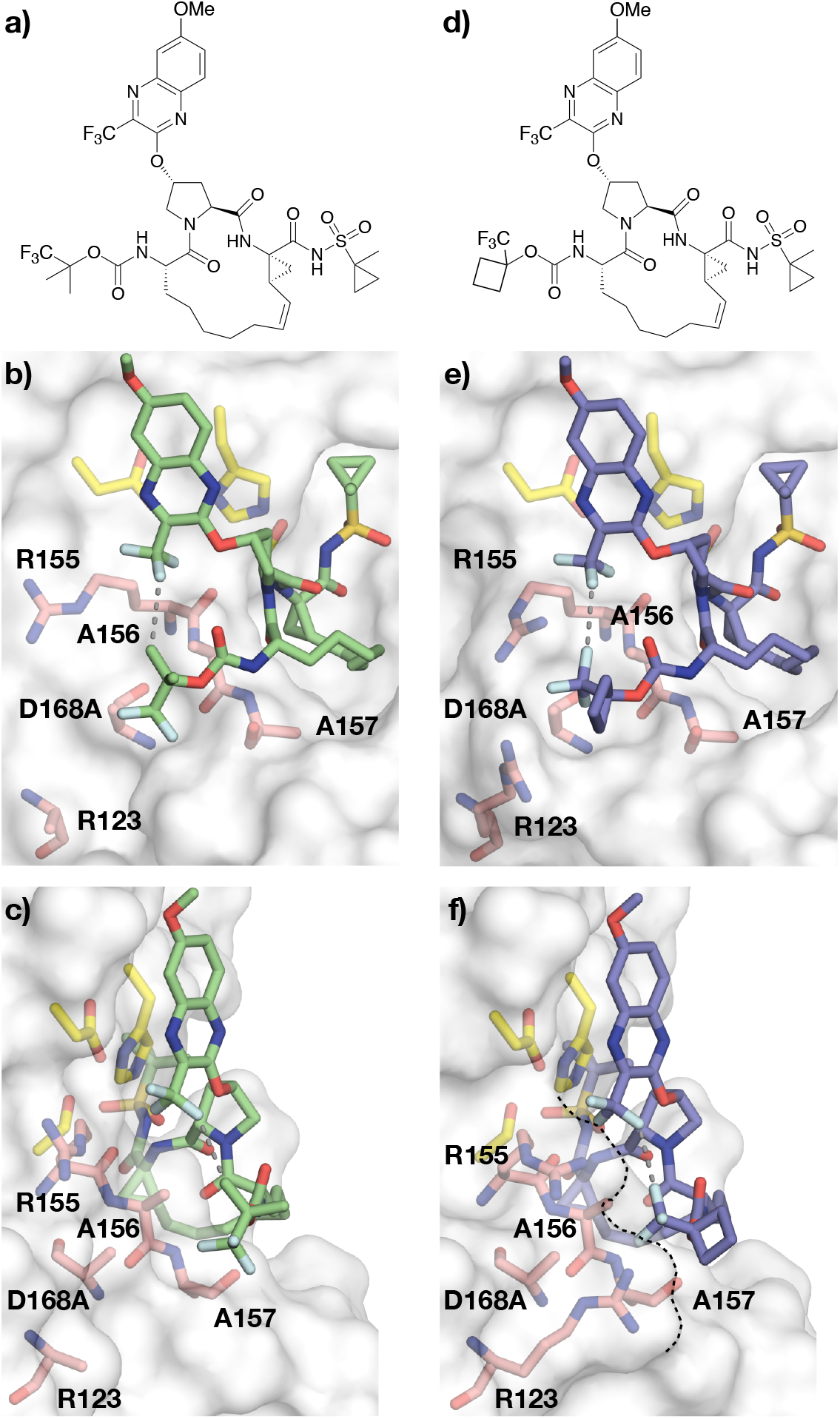
Intermolecular fluorine interaction mediated by R155. Chemical structures and cocrystal structures of **(a-c) 8F** (PDB ID: 7L7L) and **(d-f) 9F** (PDB ID: 7L7N) bound to the D168A protease variant, front **(b** and **e)** and side **(c** and **f)** views. Residues of the S4 pocket and around the fluorine atoms are labeled. The protease is shown as a surface representation. The dotted black line outlined the complementary electrostatic pocket formed by R155 and R123. The dotted gray lines represent the closest distance between the CF_3_ substituent on the quinoxaline and the P4 capping group of **8F** (3.3 Å) and **9F** (2.9 Å).

Evidence of interdependence between the binding mode of the P2^+^ and P4 cap groups is also present when analyzing the binding conformations of **9F** and **4F** bound to the D168A protease variant. The overall binding mode of these two PIs are similar; however, the orientation of the 1-(trifluoromethyl) cyclobutyl P4 cap group is reversed. The binding mode of **4F** orients the P4 cap trifluoromethyl moiety toward R123, while **9F** binds with that moiety contacting the P2^+^ quinoxaline trifluoromethyl substituent. Surprisingly, the two trifluoromethyl moieties of **9F** are 2.9 Å apart despite being an unfavorable dipole-dipole interaction. To stabilize this binding conformation, R155 adopts a unique rotamer that positions the guanidinium moiety between these two trifluoromethyl groups. R155 thus shields the dipole-dipole repulsion force and bridges an extensive electrostatic network with the 3-trifluoromethyl of the quinoxaline, the 1-trifluoromethyl of the P4 capping group, and the guanidinium side chain of R123. Therefore, the co-crystal structure of **9F** bound to the D168A protease variant shows R155 serving as the mediator between the trifluoromethyl groups and together with **4F** provides an example of interdependence between the P2^+^ and P4 cap groups that can be leveraged to deviate from the P2-P4 macrocyclic scaffolds of the current drugs while retaining potency and potentially avoiding RASs at A156.

## DISCUSSION

Fluorination has been used in drug design to modulate physiochemical properties and bioactivity of small molecules and to target resistant variants (10, 12, 14). In this study, we elucidated the molecular mechanisms by which fluorination can serve as a general strategy to combat drug resistance. Specifically, we explained the molecular mechanism by which PIs with fluorinated P4 caps retain better potency against the resistant D168A protease variant compared to non-fluorinated analogues. The D168A RAS disrupts the electrostatic network involving R155 and R123 which provides a stable hydrophobic surface for ligand binding (32, 33). Although all PIs lost potency against the D168A protease variant, fluorine substitutions in P4 caps enabled retaining better potency compared to non-fluorinated analogues often through conformational rearrangement.

The crystal structures determined revealed that fluorine substitutions often result in a change in inhibitor binding conformation, with most fluorinated PIs sampling two conformations. Comparative structural analysis of non-fluorinated and fluorinated PIs bound to both WT and D168A protease variant revealed that fluorination of the P4 caps allows the PIs to better adapt to the structural and electrostatic changes in the S4 pocket resulting from the D168A RAS. In the D168A variant, interactions of the fluorine atoms stabilize the S4 pocket and decrease conformational dynamics of R123 through orthogonal multipolar and electrostatic interactions. The fluorinated P4 caps better fill the space created by the Asp to Ala mutation through adopting varied conformations and enhancing vdW contacts. The interaction between fluorine and alanine is favorable, as fluorine has hydrophobic characteristics. In contrast in the WT protease, the fluorine atoms of the P4 caps avoid electrostatic repulsion with D168, which can form a salt bridge with R123, causing the fluorine atoms to be solvent exposed in most cases. The D168A RAS, however, permits the fluorine atoms to form favorable interactions with R123, better fill in the S4 pocket –which is further aided by the ∼18% larger vdW radius of fluorine compared to hydrogen (1.20 Å versus 1.47 Å)– and eliminates the potential of electrostatic repulsion (34). Altogether, PIs with fluorinated P4 caps avoid repulsive interaction with D168 in WT protease and adopt an alternate binding mode in the D168A resistant variant to optimize interactions.

Our results indicate that design of fluorine substitutions to small molecule inhibitors needs to account for conformational changes that can be induced by fluorination. A computational algorithm, FMAP, targets backbone carbonyls and aims to identify potential fluorination sites to participate in orthogonal multipolar interactions but does not account for conformational changes resulting from fluorination (35). Here, we observed that fluorination is almost always associated with a change in inhibitor binding conformation. In addition, we describe orthogonal multipolar interaction between fluorine and the imine carbon of arginine, and a potential repulsive force with the carboxylic acid side chain of aspartate (36).

The latest drugs, GLE and VOX, have successfully incorporated fluorine into the P2-P4 macrocyclic scaffold to target all genotypes with pico-molar potencies driven in part by fluorine specific interactions, including three orthogonal multipolar interactions, and a “caged” fluorine-induced hydrogen bond that aids in pre-organizing the inhibitor for binding (**Fig. 2 and Fig. S14**) (6, 8). However, both GLE and VOX are highly susceptible to RAS at A156, thereby enabling the possibility of cross-resistance (37-39). The P2-P4 macrocycle is currently the Achilles’ heel of all three FDA-approved drugs, as RASs at position 156 cause steric clash with the P2-P4 macrocycle (4, 5, 7). Scaffold diversification including optimized P1-P3 macrocycles, and fluorination may be needed to rationally design novel inhibitors to mitigate resistance should future RASs render all FDA-approved HCV NS3/4A PIs ineffective. The intramolecular interaction between the P2^+^ and P4 moieties of **1F** reveals a potential strategy to target RASs occurring at A156 by further stabilizing the binding mode without a P2-P4 macrocycle.

In summary, here we show that strategic incorporation of fluorine atoms proximal to the sites of RASs in the HCV NS3/4A protease resulted in PIs with improved resistance profiles. P4 cap fluorination allowed PIs to adapt to the RAS-induced structural and electrostatic changes in the S4 pocket of the protease by sampling alternate, more favorable, binding conformations. Thus, we present a novel molecular strategy in inhibitor design to combat drug resistance.

## METHODS

### Inhibitor design and synthesis

The compounds were computationally modeled using Maestro from Shrodinger starting from the crystal structure of parent compound bound to WT protease (PDB ID: 5VOJ). The inhibitors were synthesized in-house using our convergent reaction sequence previously reported with minor modifications (40).

### Expression and purification of NS3/4A constructs

The HCV GT1a NS3/4A protease gene described in the Bristol Myers Squibb patent was synthesized by GenScript and cloned into a PET28a expression vector (41). Cys159 was mutated to a serine residue to prevent disulfide bond formation and facilitate crystallization. The D168A gene was engineered using the site-directed mutagenesis protocol from Stratagene. Protein expression and purification were carried out as previously described (42). Briefly, transformed *Escherichia coli* BL21(DE3) cells were grown in TB media containing 30 μg/mL of kanamycin antibiotic at 37 °C. After reaching an OD_600_ of 0.7, cultures were induced with 1 mM IPTG and harvested after 3 h of expression. Cells were pelleted by centrifugation, resuspended in resuspension buffer (RB) [50 mM phosphate buffer, 500 mM NaCl, 10% glycerol, 2 mM β-ME, pH 7.5] and frozen at −80 °C for storage.

Cell pellets were thawed and lysed via cell disruptor (Microfluidics Inc.) two times to ensure sufficient DNA shearing. Lysate was centrifuged at 19,000 rpm, for 25 min at 4 °C. The soluble fraction was applied to a nickel column (Qiagen) pre-equilibrated with RB. The beads and soluble fraction were incubated at 4 °C for 1.5 h and the lysate was allowed to flow through. Beads were washed with RB supplemented with 20 mM imidazole and eluted with RB supplemented with 200 mM imidazole. The eluent was dialyzed overnight (MWCO 10 kD) to remove the imidazole, and the His-tag was simultaneously removed with thrombin treatment. The eluate was judged >90% pure by polyacrylamide gel electrophoresis, concentrated, flash frozen, and stored at −80 °C.

### Correction for the inner filter effect

The inner filter effect (IFE) for the NS3/4A protease substrate was determined using a previously described method (43). Briefly, fluorescence end-point readings were taken for substrate concentrations between 0 μM and 20 μM. Afterward, free 5-FAM fluorophore was added to a final concentration of 25 μM to each substrate concentration and a second round of fluorescence end-point readings was taken. The fluorescence of free 5-FAM was determined by subtracting the first fluorescence end point reading from the second round of readings. IFE corrections were then calculated by dividing the free 5-FAM florescence at each substrate concentration by the free 5-FAM florescence at zero substrate.

### Determination of Michaelis–Menten (K_m_) constant

K_m_ constants for GT1 and D168A protease were previously determined (40). Briefly, a 20 μM concentration of substrate [Ac-DE-Dap(QXL520)-EE-Abu-γ-[COO]AS-C(5-FAMsp)-NH2] (AnaSpec) was serially diluted into assay buffer [50 mM Tris, 5% glycerol, 10 mM DTT, 0.6 mM LDAO, and 4% dimethyl sulfoxide] and proteolysis was initiated by rapid injection of 10 μL protease (final concentration 20 nM) in a reaction volume of 60 μL. The fluorescence output from the substrate cleavage product was measured kinetically using an EnVision plate reader (Perkin-Elmer) with excitation wavelength at 485 nm and emission at 530 nm. Inner filter effect corrections were applied to the initial velocities (*V*_o_) at each substrate concentration. *V*_o_ versus substrate concentration graphs were globally fit to the Michaelis–Menten equation to obtain the K_m_ value.

### Enzyme inhibition assays

For each assay, 2 nM of NS3/4A protease (GT1a and D168A) was pre-incubated at room temperature for 1 h with increasing concentration of inhibitors in assay buffer (50 mM Tris, 5% glycerol, 10 mM DTT, 0.6 mM LDAO, and 4% dimethyl sulfoxide, pH 7.5). Inhibition assays were performed in non-binding surface 96-well black half-area plates (Corning) in a reaction volume of 60 μL. The proteolytic reaction was initiated by the injection of 5 μL of HCV NS3/4A protease substrate (AnaSpec), to a final concentration of 200 nM and kinetically monitored using a Perkin Elmer EnVision plate reader (excitation at 485 nm, emission at 530 nm). Three independent data sets were collected for each inhibitor with each protease construct. Each inhibitor titration included at least 12 inhibitor concentration points, which were globally fit to the Morrison equation to obtain the *K*_i_ value.

### Crystallization and structure determination

Protein expression and purification were carried out as previously described with slight modifications (42). Briefly, the Histrap purified WT1a or D168A mutant protein was thawed, concentrated to 3 mg/mL, and loaded on a HiLoad Superdex75 16/60 column equilibrated with gel filtration buffer (25 mM MES, 500 mM NaCl, 10% glycerol, and 2 mM DTT, pH 6.5). The protease fractions were pooled and concentrated to 25 mg/mL with an Amicon Ultra-15 10 kDa filter unit (Millipore). The concentrated samples were incubated for 1 h with 3:1 or 6:1 molar excess of inhibitor. Diffraction-quality crystals were obtained within 2 days by mixing equal volumes of concentrated protein solution with precipitant solution (20–30% PEG-3350, 0.1 M sodium MES buffer, 1-7% ammonium sulfate, pH 6.5) at RT or 15 °C in 24-well VDX hanging drop trays. Crystals were harvested and data was collected at 100 K. Cryogenic conditions contained the precipitant solution supplemented with 15% glycerol or ethylene glycol.

X-ray diffraction data were collected in-house using our Rigaku X-ray system with a Saturn 944 detector. All datasets were processed using HKL-3000 (44). Structures were solved by molecular replacement using PHASER (45). Model building and refinement were performed using Coot (46) and PHENIX (47), respectively. The final structures were evaluated with MolProbity (48) prior to deposition in the PDB. To limit the possibility of model bias throughout the refinement process, 5% of the data were reserved for the free R-value calculation (49). Structure analysis, superposition and figure generation were done using PyMOL (50). X-ray data collection and crystallographic refinement statistics are presented in **Table S1**. Crystals of **4F** bound to the WT NS3/4A protease did not contain adequate electron density to accurately build in the entire inhibitor, but the partial electron density of the ligand guided the modeled structure.

### Structural analysis

Superpositions were performed in PyMol using the Cα atoms of active site residues 137-139 and 154-160 of the protease. The D168A-parent compound complex structures were used as references for the alignments in sets 1-7. The van der Waals contact energies between the protease and the inhibitors were computed using a simplified Lennard-Jones potential as described previously (51).

### Molecular modeling

Modeling of **4F** bound to the HCV WT protease was done in Maestro. The cocrystal structures of **4** bound to the WT protease and **4F** bound to the D168A were used as starting models. Chain A of the complex structure of **4F** bound to the D168A variant was superimposed on the structure of **4** bound to the WT protease (RMSD 0.9 Å). The mutate function in Maestro was used to mutate Ala to Asp at the 168 position. The rotamer function was used to select an Asp conformation similar to what is seen in the **4**-WT structure. The newly generated Asp 168 residue was then minimized.

Modeling of the WT and D168A complex structures with **2** followed a similar procedure as above. More specifically, staring with the structures of **2F**_***R***_ bound to WT and D168A, the fluorine atoms of CF_3_ group were replaced with hydrogens in Maestro to generate the complex structure with **2**. The complex model structures were then minimized.

### Protein Data Bank accession number

The atomic coordinates have been deposited in the RCSB Protein Data Bank. The accession codes for the WT HCV protease – inhibitor complex structures are; **3**, 7MM4; **4**, 7MM2; **5**, 7MM3; **1F**, 6DIS; **2F**_***S***_, 7MM7; **2F**_***R***_, 7MMA; **3F**, 7MM6; **5F**, 7MM9; **6F**, 7MM5; and **7F**, 7MM8. The accession codes for the D168A variant of the HCV protease – inhibitor complex structures are; **1**, 7MM6; **3**, 7MMC; **5**, 7MMB; **7**, 7MMD; **1F**, 6DIW; **2F**_***S***_, 7MMI; **2F**_***R***_, 7MML; **3F**, 7MMH; **4F**, 7MMG; **5F**, 7MMK; **6F**, 7MMF; and **7F**, 7MMJ.

## Acknowledgements

This research used resources of the Advanced Photon Source, a U.S. Department of Energy (DOE) Office of Science User Facility operated for the DOE Office of Science by Argonne National Laboratory under Contract No. DE-AC02-06CH11357. We thank beam line specialist at 23-ID-B for their help in data collection. This work was supported by the National Institute of Allergy and Infectious Diseases (R01 AI085051) and National Institute of General Medical Sciences (R01 GM135919). JZ and ANM were also supported by the National Institute of General Medical Sciences of the NIH (F31 GM119345 and F31 GM131635 respectively).

## TOC Graphic

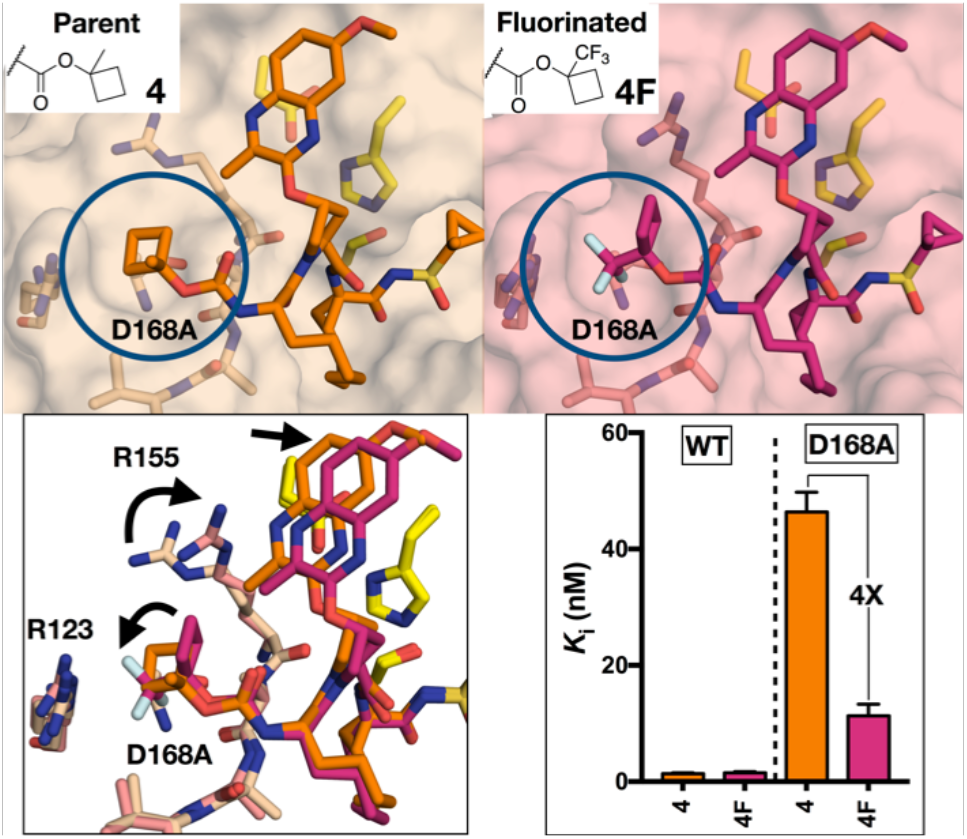

