## Supplemental information for "Deciphering the Molecular Mechanism of HCV Protease Inhibitor Fluorination as a General Approach to Avoid Drug Resistance"

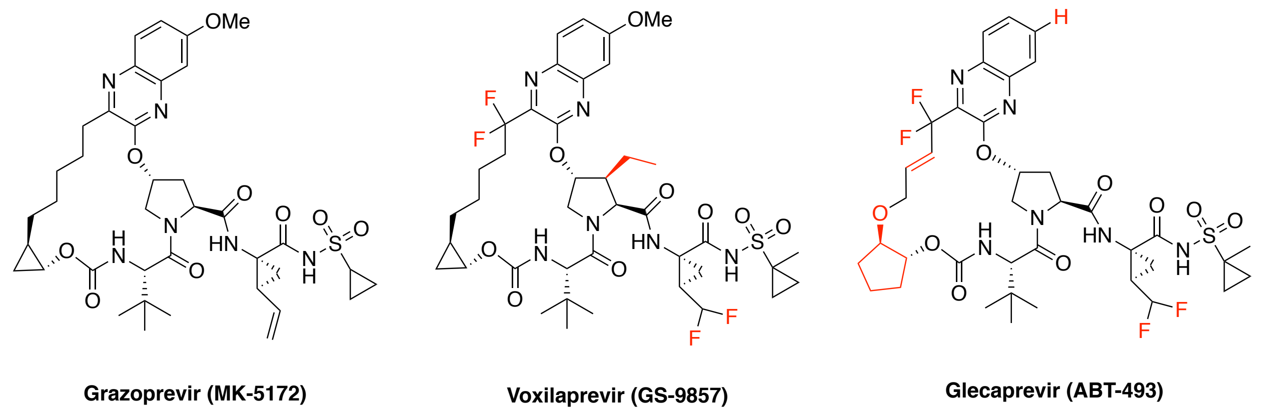


**Figure S1. FDA-Approved HCV NS3/4A Protease Inhibitors Currently in Clinic**
The atoms highlighted red in voxilaprevir (VOX) and glecaprevir (GLE) indicate chemical differences compared to grazoprevir (GZR).

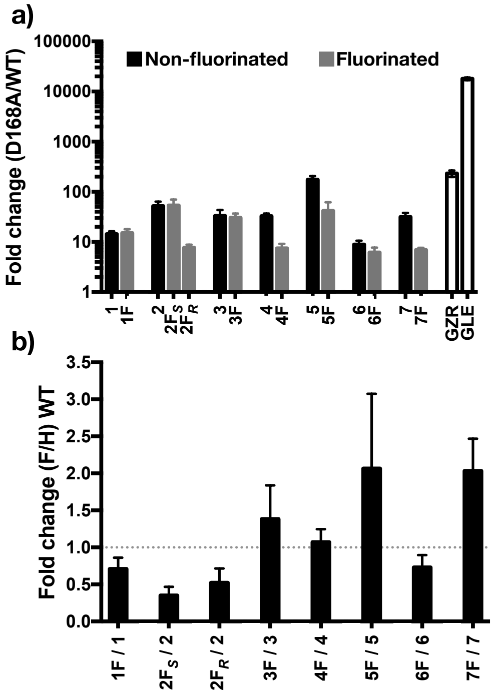


**Figure S2. Fold change in inhibitor potency for fluorinated inhibitors and their non-fluorinated analogs**. **(a)** Fold change in inhibitor potencies (enzyme inhibition constants) resulting from the D168A RAS. **(b)** Fold change in potency against the WT HCV NS3/4A protease resulting from P4 cap fluorination.

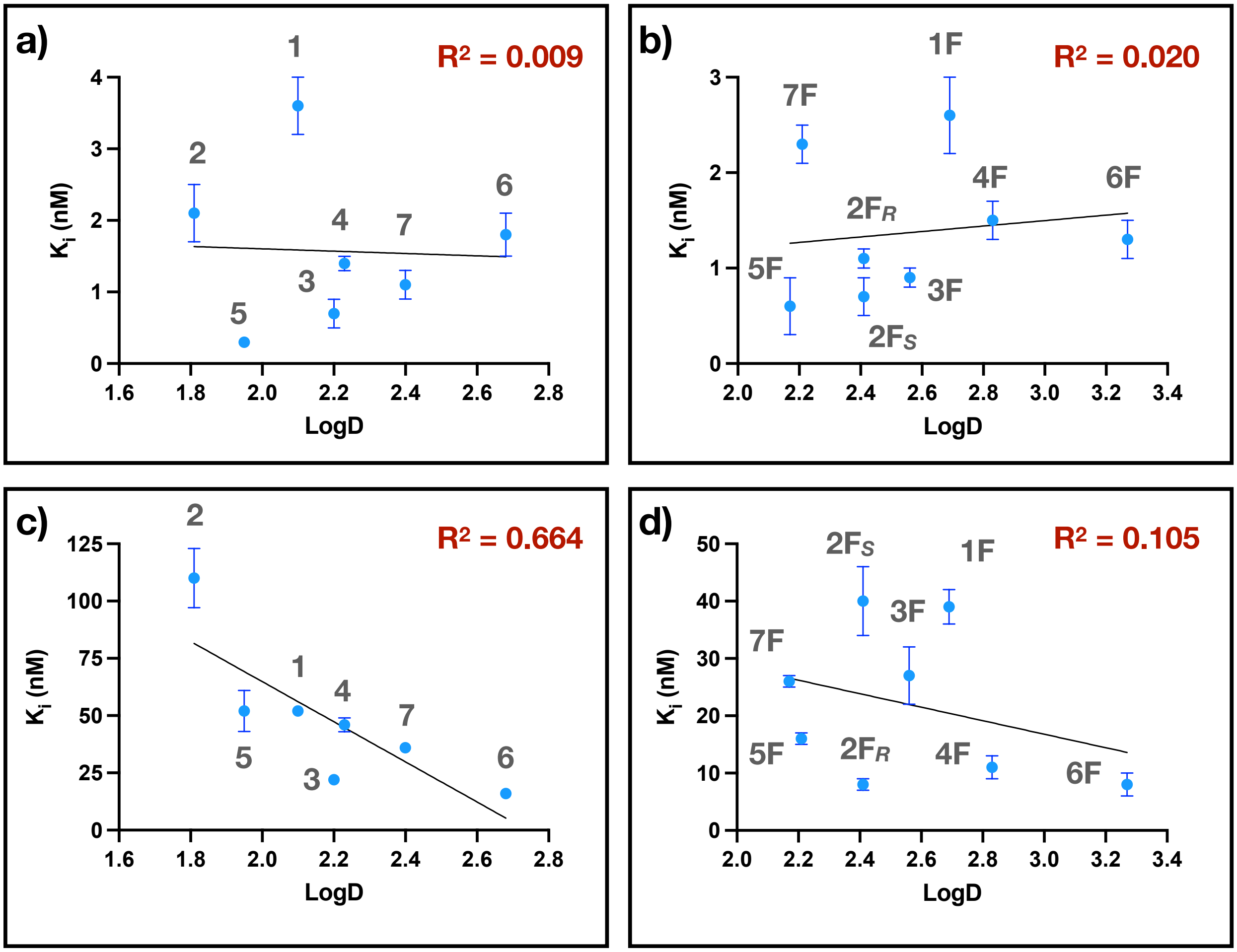


**Figure S3. Lipophilicity does not explain the increase in potency of the fluorinated inhibitors against the D168A protease variant**

The correlation of inhibition constants against the WT HCV protease and LogD for (**a**) non-fluorinated and (**b**) fluorinated inhibitors. The correlation of inhibition constants against the D168A HCV protease and LogD for (**c**) non-fluorinated and (**d**) fluorinated inhibitors. The logD was calculated using Marvin Sketch.

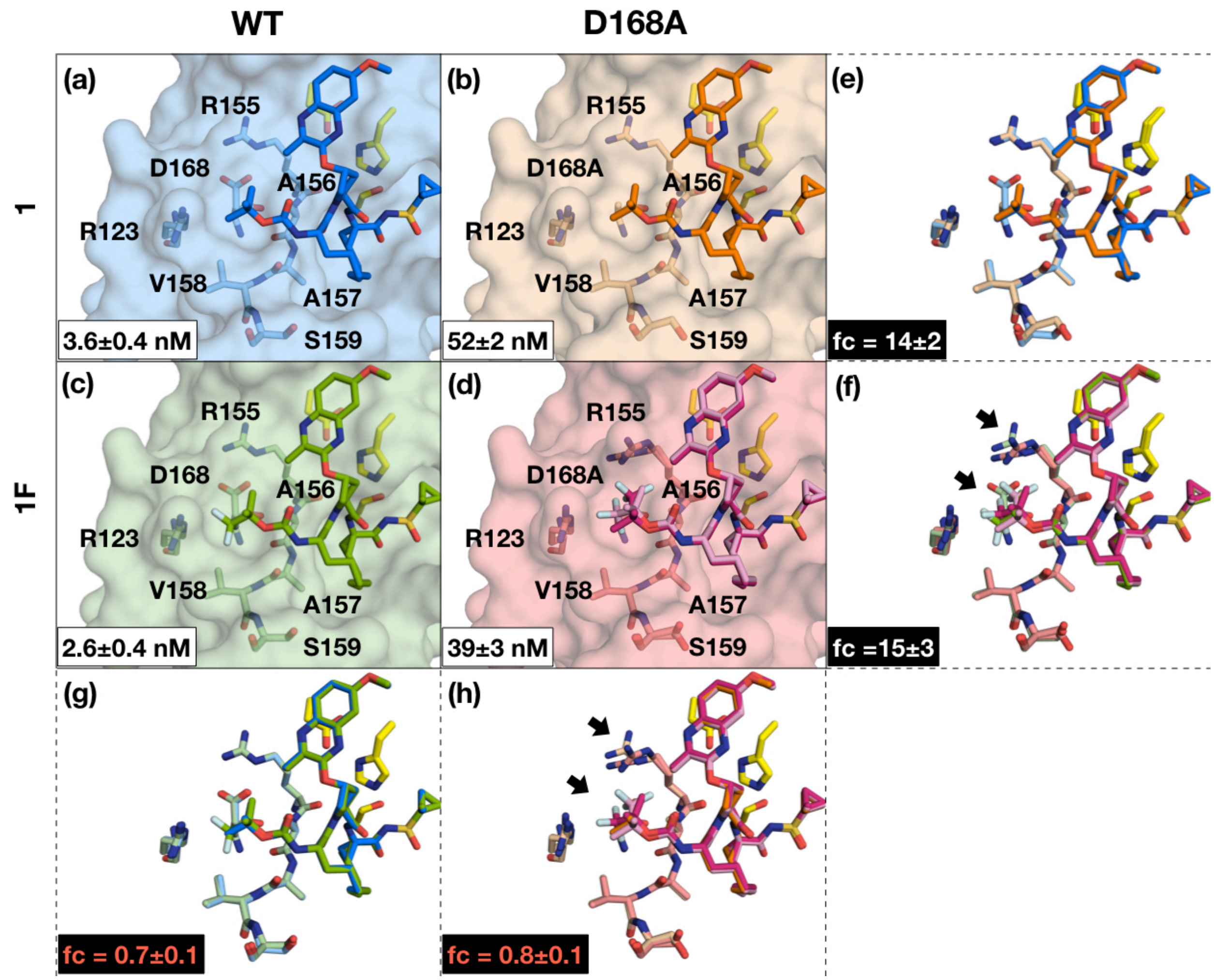


**Figure S3.** Co-crystal structures of **1** bound to HCV NS3/4A protease **(a)** WT (GT-1a) and **(b)** D168A, and **1F** bound to **(c)** WT and **(d)** D168A. The binding affinity for each complex is indicated within the inset in panels **(a-d)**. Structural differences for binding to WT versus D168A are shown by superpositions for **(e)** **1** and **(f)** **1F**; and binding of **1** versus **1F** to **(g)** WT and **(h)** D168A protease, respectively. The fold change (fc) in binding affinity is indicated in panels **(e and f)** relative to WT, and relative to the parent inhibitor in **(g)** and **(h)** for WT and D168A, respectively. The inhibitors as well as residues of the catalytic triad and S4 pocket (labeled) are shown as sticks. Inhibitors with two conformations are shown in two shades of the same color, where the darker shade represent the highest occupancy conformation. Arrows in panels **f-h** indicate structural differences between the superimposed structures.

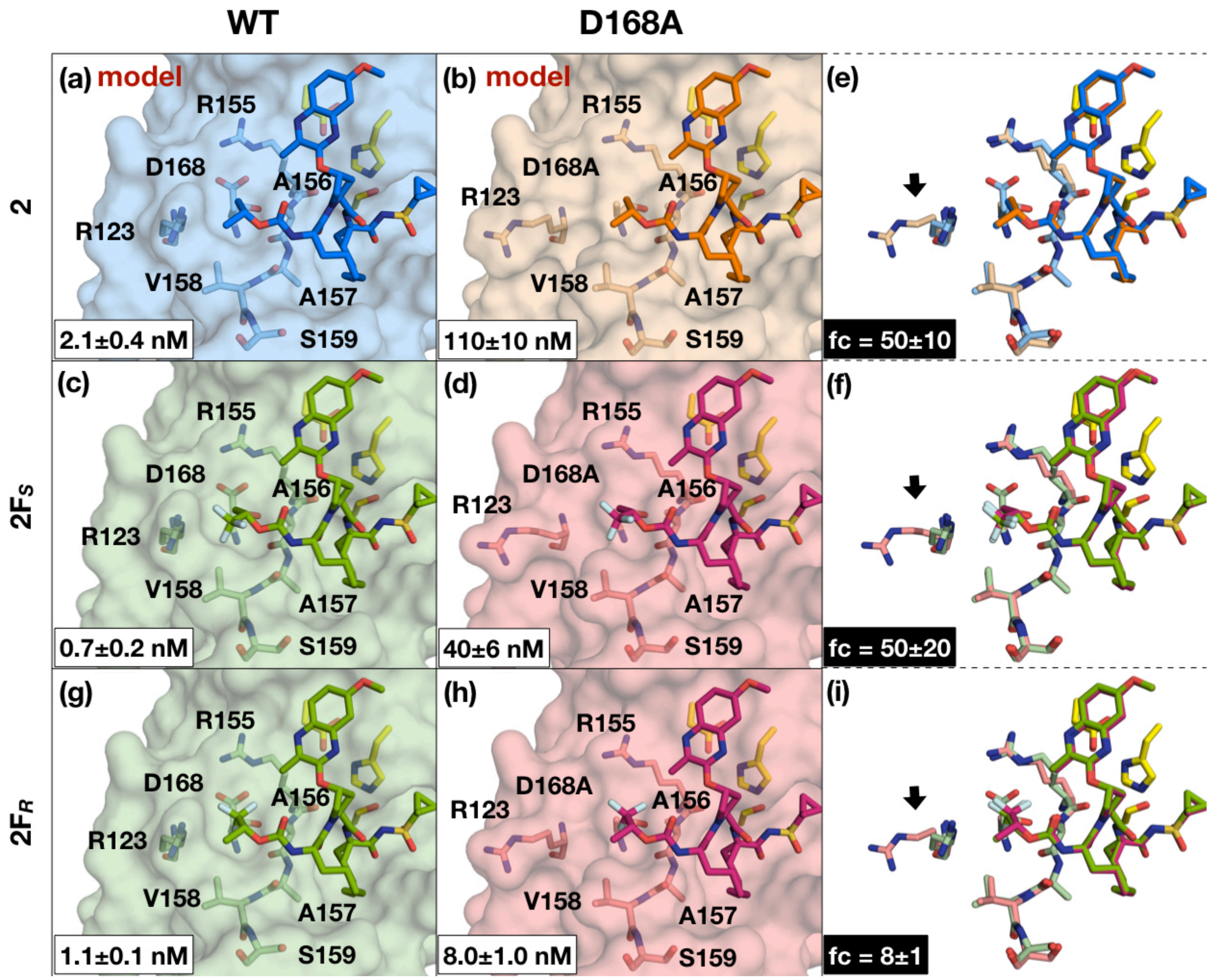


**Figure S4.** Structural models of **2** bound to HCV NS3/4A protease **(a)** WT (GT-1a) and **(b)** D168A; and cocrystal structures of **2F*S*** bound to **(c)** WT and **(d)** D168A, of **2F*R*** bound to **(g)** WT and **(h)** D168A. The binding affinity for each complex is indicated within the inset in panels **(a-h)**. Structural differences for binding to WT versus D168A are shown by superpositions for **(e) 2, (f) 2F*S* (i) 2F*R***, where the arrows indicate structural differences. The fold change (fc) in binding affinity is indicated in panels **(e, f and i)** relative to WT. The inhibitors as well as residues of the catalytic triad and S4 pocket (labeled) are shown as sticks. Inhibitors with two conformations are shown in two shades of the same color, where the darker shade represent the highest occupancy conformation.

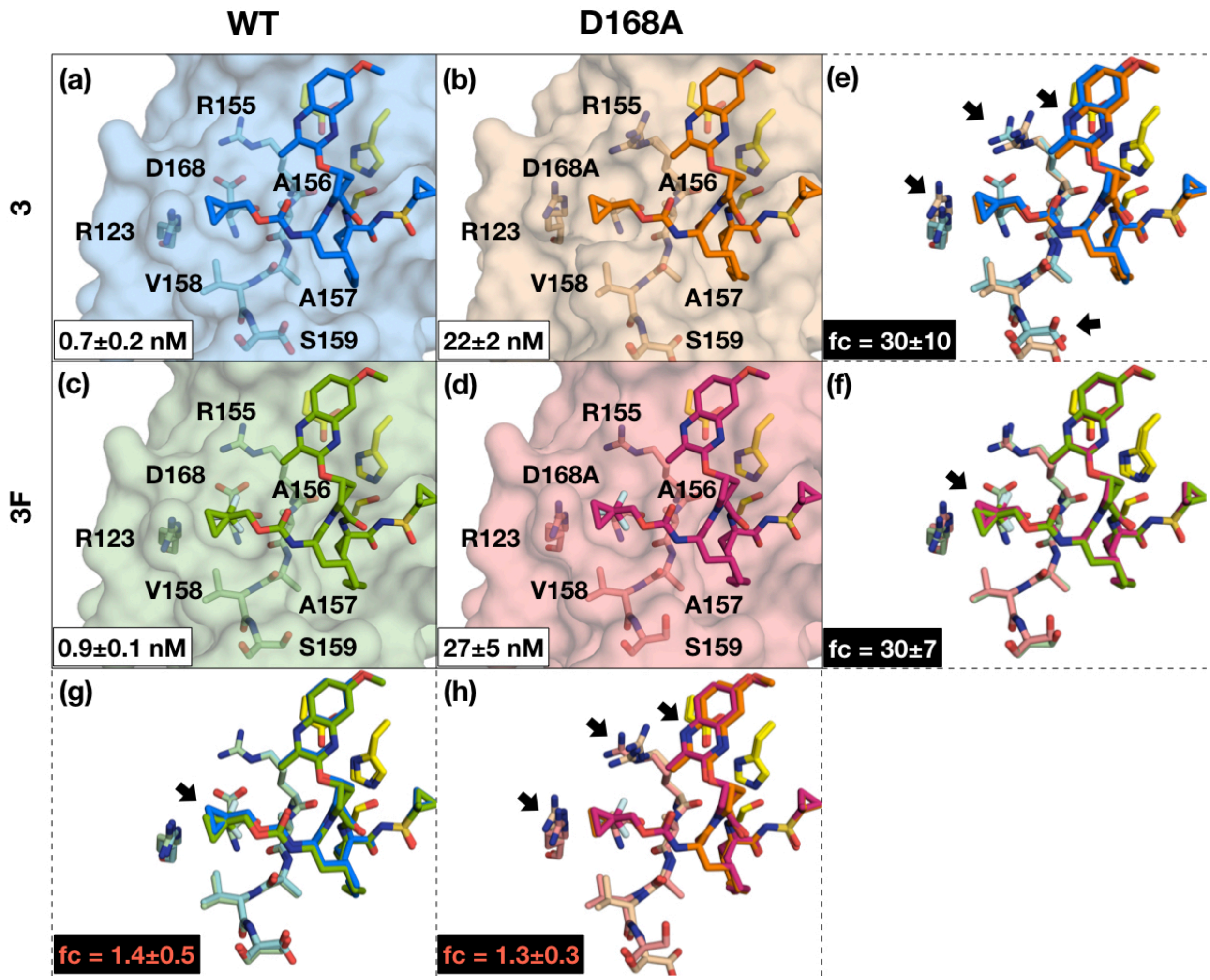


**Figure S5.** Co-crystal structures of **3** bound to HCV NS3/4A protease **(a)** WT (GT-1a) and **(b)** D168A, and **3F** bound to **(c)** WT and **(d)** D168A. The binding affinity for each complex is indicated within the inset in panels **(a-d)**. Structural differences for binding to WT versus D168A are shown by superpositions for **(e) 3** and **(f) 3F**; and binding of **3** versus **3F** to **(g)** WT and **(h)** D168A protease, respectively. The fold change (fc) in binding affinity is indicated in panels **(e and f)** relative to WT, and relative to the parent inhibitor in **(g)** and **(h)** for WT and D168A, respectively. The inhibitors as well as residues of the catalytic triad and S4 pocket (labeled) are shown as sticks. Inhibitors with two conformations are shown in two shades of the same color, where the darker shade represent the highest occupancy conformation. Arrows in panels **e**-**h** indicate structural differences between the superimposed structures.

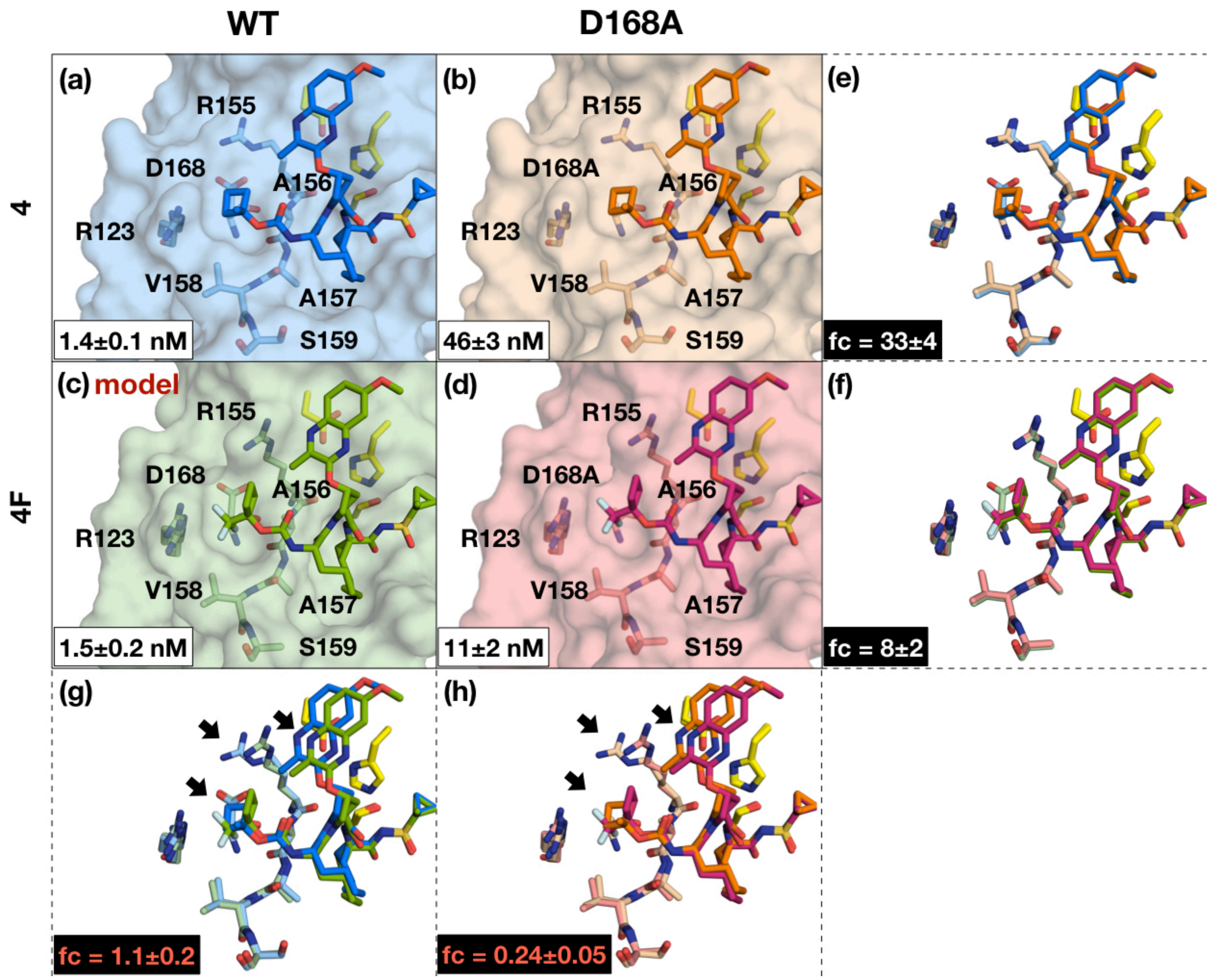


**Figure S6.** Co-crystal structures of **4** bound to HCV NS3/4A protease **(a)** WT (GT-1a) and **(b)** D168A, and **4F** bound to **(c)** WT and **(d)** D168A. The binding affinity for each complex is indicated within the inset in panels **(a-d)**. Structural differences for binding to WT versus D168A are shown by superpositions for **(e) 4** and **(f) 4F**; and binding of **4** versus **4F** to **(g)** WT and **(h)** D168A protease, respectively. The fold change (fc) in binding affinity is indicated in panels **(e** and **f)** relative to WT, and relative to the parent inhibitor in **(g)** and **(h)** for WT and D168A, respectively. The inhibitors as well as residues of the catalytic triad and S4 pocket (labeled) are shown as sticks. Inhibitors with two conformations are shown in two shades of the same color, where the darker shade represent the highest occupancy conformation. Arrows in panels **g-h** indicate structural differences between the superimposed structures.

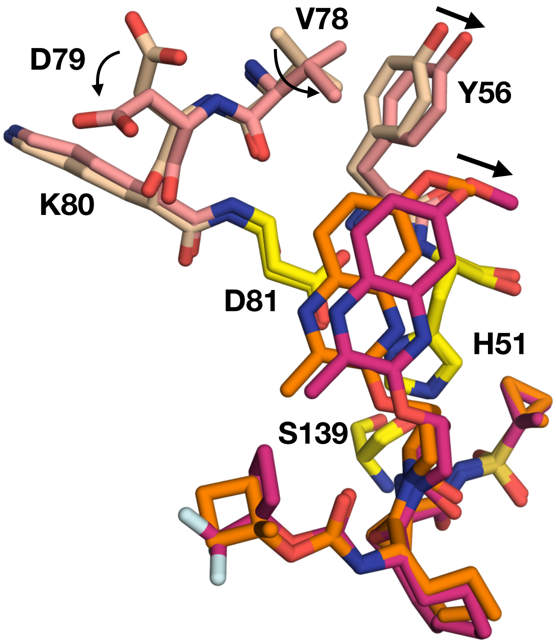


**Figure S7. Rearrangement of the S2$^{+}$ subsite and the P2$^{+}$ quinoxaline of the inhibitor with fluorination of the P4 cap.**

The residues making the S2$^{+}$subsite, the catalytic triad (yellow), and inhibitors **4** and **4F** are shown as sticks and labeled. **4** and **4F** are colored orange and salmon respectively. The arrows indicate movement of the inhibitor and protease residues.

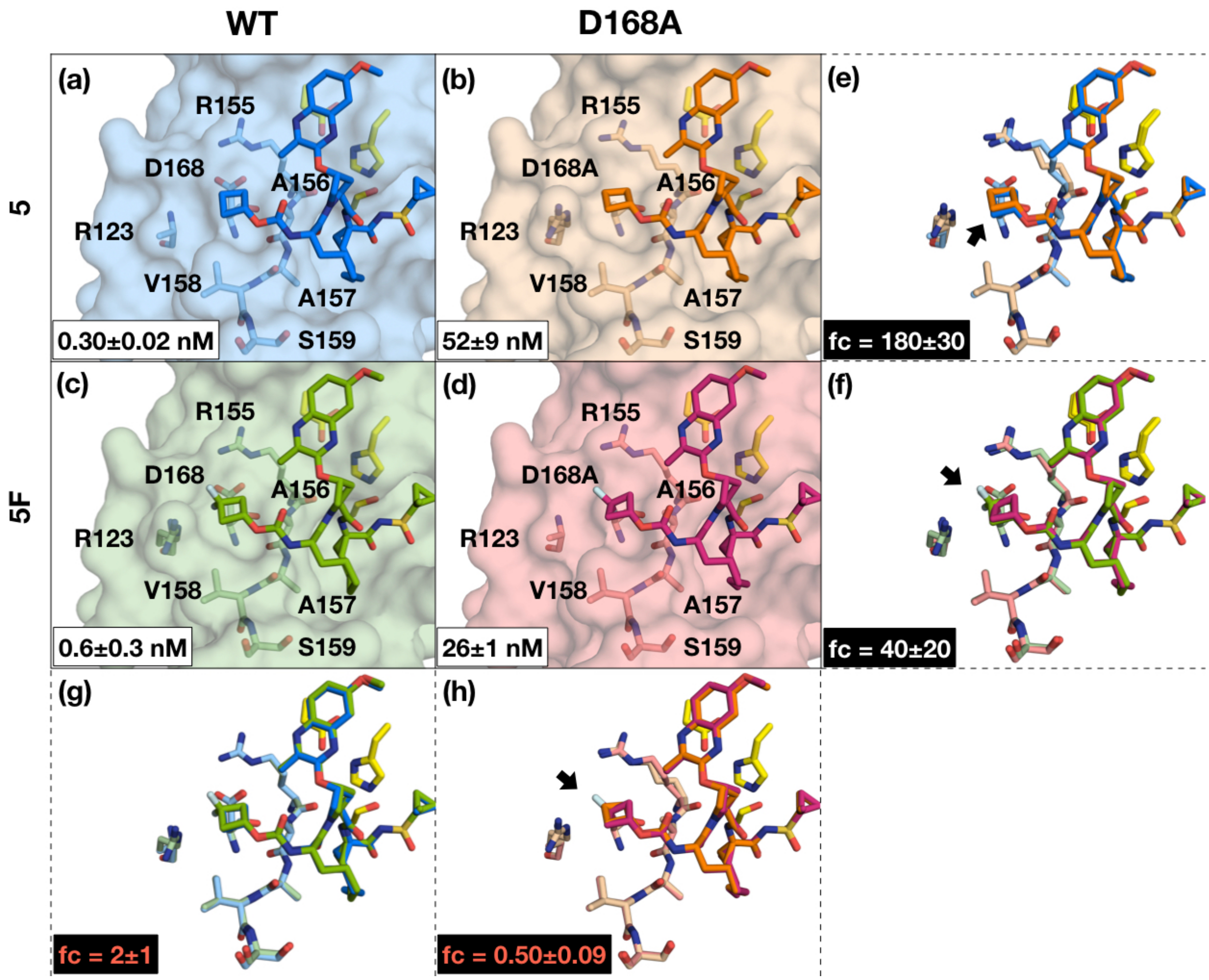


**Figure S8.** Co-crystal structures of **5** bound to HCV NS3/4A protease **(a)** WT (GT-1a) and **(b)** D168A, and **5F** bound to **(c)** WT and **(d)** D168A. The binding affinity for each complex is indicated within the inset in panels **(a-d)**. Structural differences for binding to WT versus D168A are shown by superpositions for **(e)** **5** and **(f)** **5F**; and binding of **5** versus **5F** to **(g)** WT and **(h)** D168A protease, respectively. The fold change (fc) in binding affinity is indicated in panels **(e and f)** relative to WT, and relative to the parent inhibitor in **(g)** and **(h)** for WT and D168A, respectively. The inhibitors as well as residues of the catalytic triad and S4 pocket (labeled) are shown as sticks. Inhibitors with two conformations are shown in two shades of the same color, where the darker shade represent the highest occupancy conformation. Arrows in panels **e**-**h** indicate structural differences between the superimposed structures.

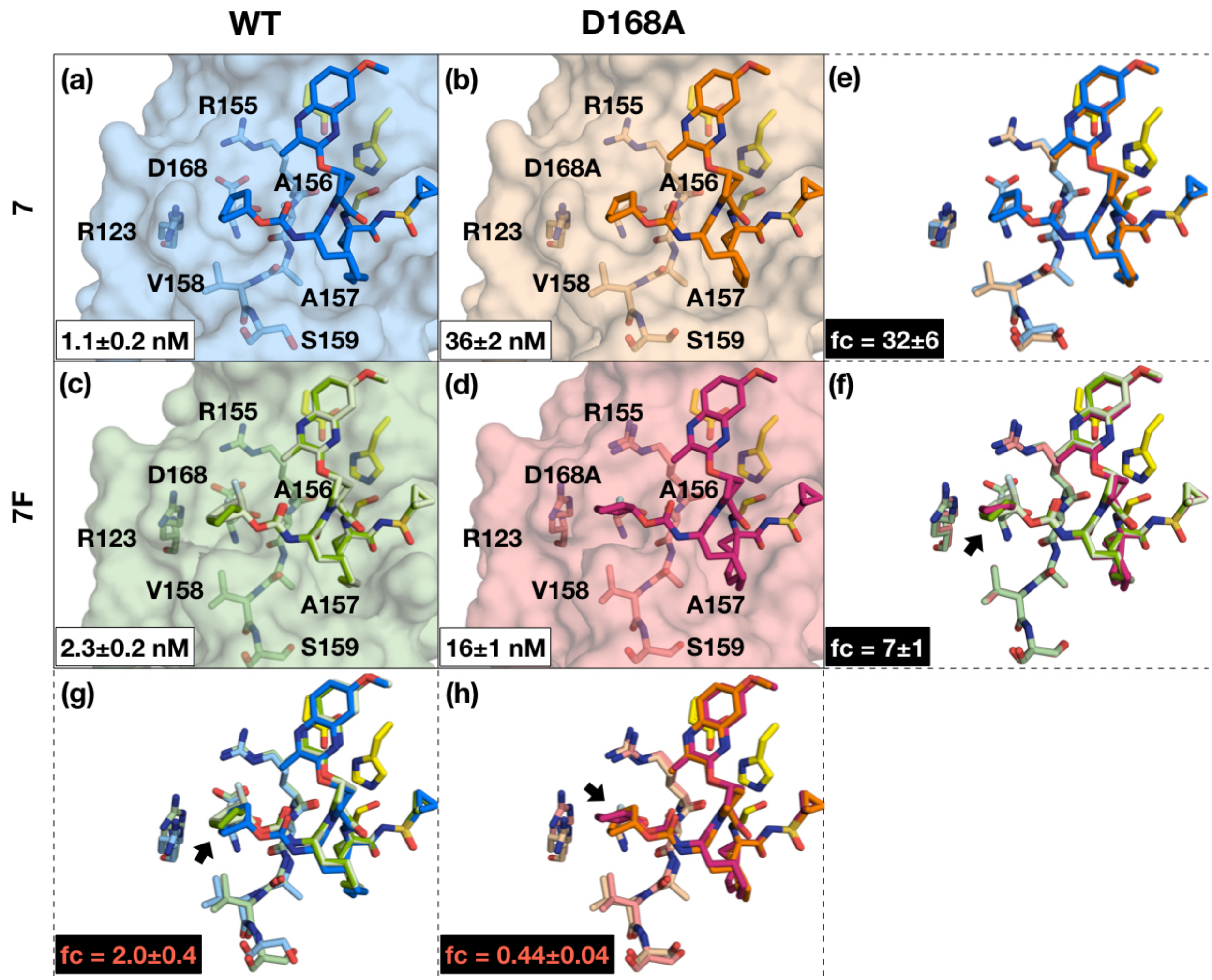


**Figure S9.** Co-crystal structures of **7** bound to HCV NS3/4A protease **(a)** WT (GT-1a) and **(b)** D168A, and **7F** bound to **(c)** WT and **(d)** D168A. The binding affinity for each complex is indicated within the inset in panels **(a-d)**. Structural differences for binding to WT versus D168A are shown by superpositions for **(e)** **7** and **(f)** **7F**; and binding of **7** versus **7F** to **(g)** WT and **(h)** D168A protease, respectively. The fold change (fc) in binding affinity is indicated in panels **(e and f)** relative to WT, and relative to the parent inhibitor in **(g)** and **(h)** for WT and D168A, respectively. The inhibitors as well as residues of the catalytic triad and S4 pocket (labeled) are shown as sticks. Inhibitors with two conformations are shown in two shades of the same color, where the darker shade represent the highest occupancy conformation. Arrows in panels **f-h** indicate structural differences between the superimposed structures.

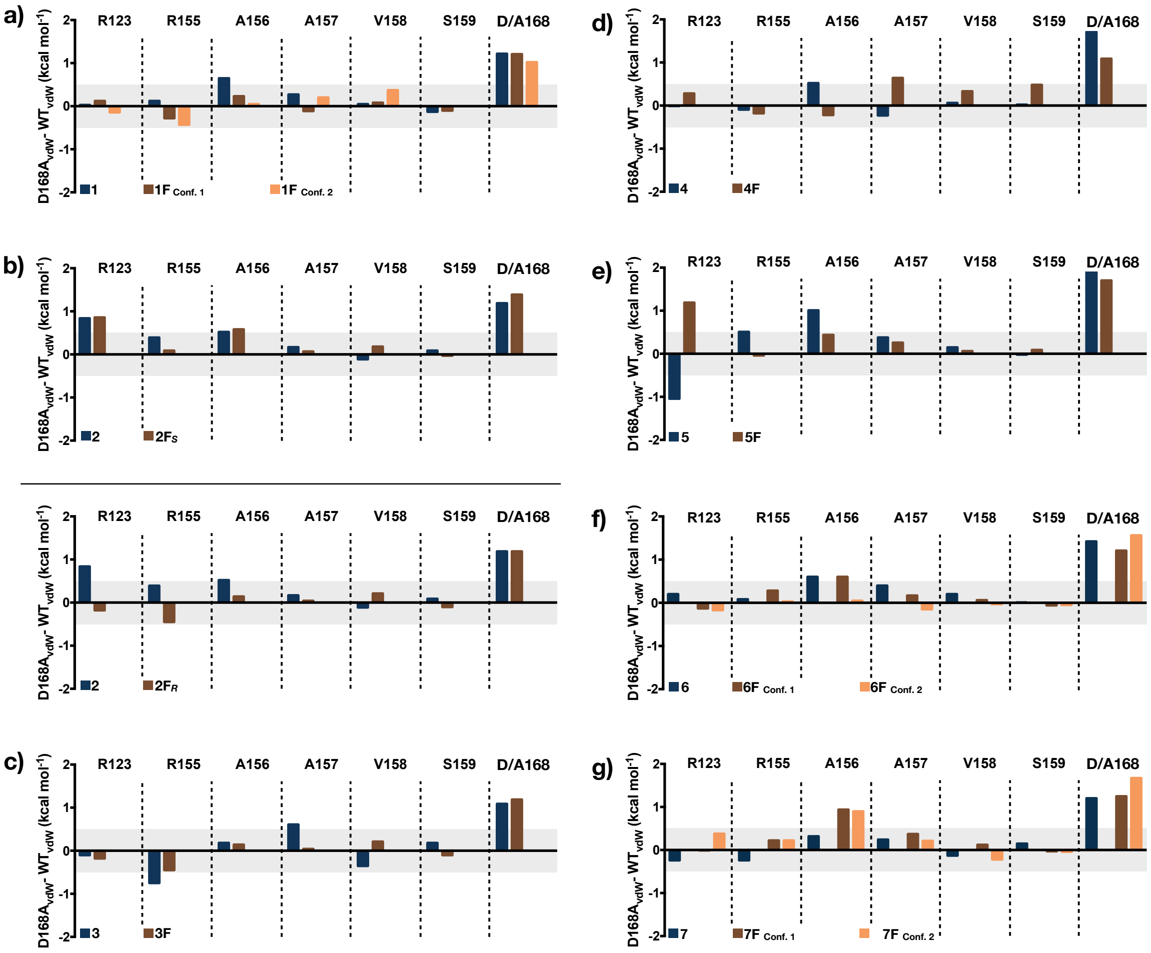


**Figure S10. Per residue van der Waals interactions of inhibitors with the protease.** Differences in vdW contacts between WT and D168A for inhibitor pairs **1-7 (a-g)** with (orange) and without (blue) fluorine substitutions for residues composing the S4 pocket. The plots on top and bottom of panel **b** are for compounds **2F*S*** and **2F*R*** respectively. Darker and lighter shades of orange indicate differences for the higher and lower occupancy conformations of the fluorinated inhibitors, respectively. Differences within 0.5 kcal mol$^{-1}$ are considered insignificant (gray bar).

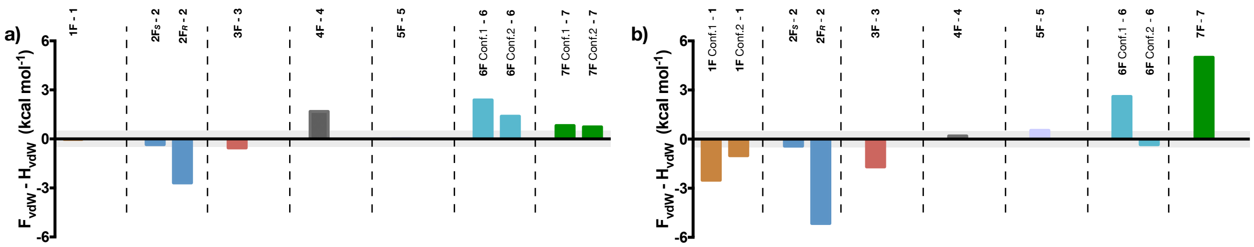


**Figure S11. Effects of inhibitor fluorination on total vdW contacts with WT (GT1a) and D168A protease.**

Differences in total vdW contacts between the inhibitor and protease in crystal structures of **(a)** WT and **(b)** D168A protease variants bound to non-fluorinated and fluorinated compounds. Differences within 0.5 kcal mol$^{-1}$ are considered insignificant (gray bar).

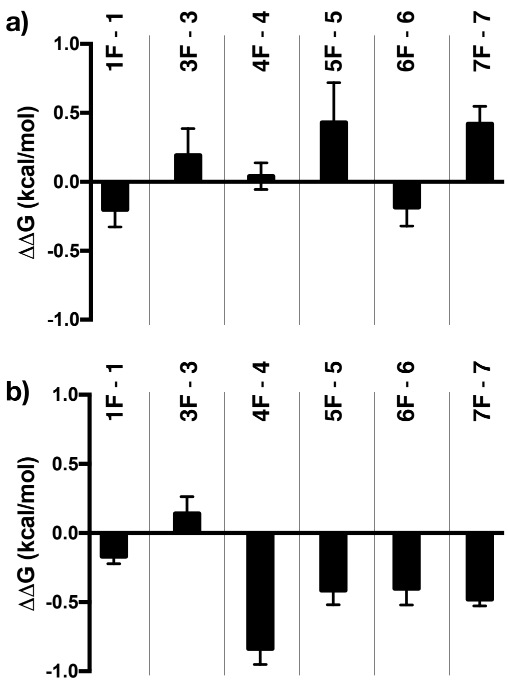


**Figure S12. Contribution of fluorination to the Gibbs free energy of binding to the (a) WT1a and (b) D168A protease variant.**

Experimental enzyme inhibition constants were used to estimate and compare the binding free energy for fluorinated versus non-fluorinated inhibitor pairs. Negative values indicate increase in potency or tighter binding due to fluorine substitutions.

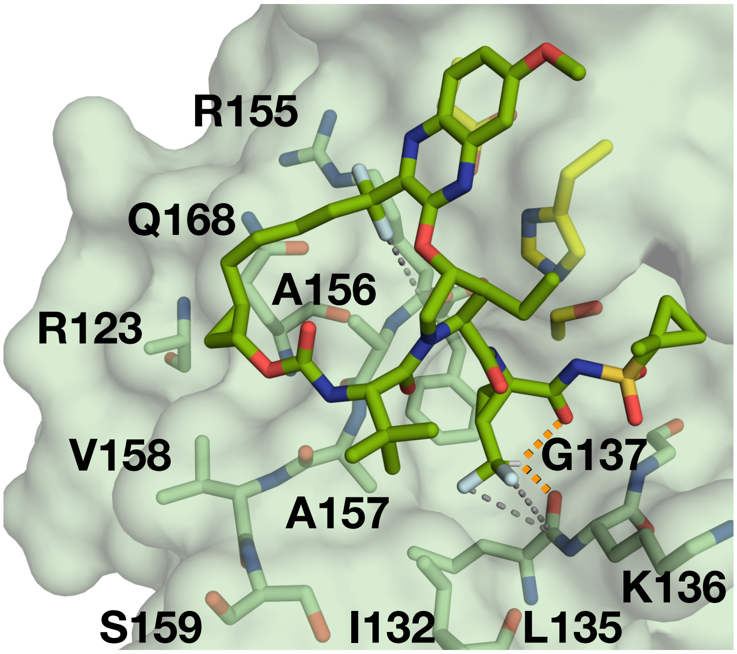


**Figure S13. Binding of voxilaprevir to the HCV NS3/4A protease**
The binding mode of voxilaprevir to the D168Q protease variant (PDB ID: 6NZT). Residues of the S4 pocket and around the fluorine atoms (light blue) are labeled; gray dotted lines represent orthogonal multipolar interactions. The orange dotted lines (2.2 and 2.6 Å) represent fluorine-induced hydrogen bonds of the P1 moiety.

**Table S1.** Inhibitory activity against wild-type (WT) GT1a HCV NS3/4A and D168A proteases in enzymatic assays, with fold change relative to WT indicated in parentheses. The error reported is from the global fit of at least three replicates.

| **Inhibitor** | **$K_i$ (nM) (Fold Change)** | |
|---|---|---|
| | **GT1a WT** | **D168A** |
| **Grazoprevir (GZR)** | 0.21 ± 0.03† | 49 ± 2† (234) |
| **1** | 3.6 ± 0.4† | 52 ± 2§ (14) |
| **2** | 2.1 ± 0.4 | 110 ± 13 (52) |
| **3** | 0.7 ± 0.2§ | 22 ± 2 (33) |
| **4** | 1.4 ± 0.1§ | 46 ± 3§ (33) |
| **5** | 0.30 ± 0.02§ | 52 ± 9§ (175) |
| **6** | 1.8 ± 0.3† | 16 ± 1† (9.0) |
| **7** | 1.1 ± 0.2§ | 36 ± 2§ (32) |
| **1F** | 2.6 ± 0.4§ | 39 ± 3§ (15) |
| **2F*S*** | 0.7 ± 0.2§ | 40 ± 6§ (54) |
| **2F*R*** | 1.1 ± 0.1§ | 8 ± 1§ (8.0) |
| **3F** | 0.9 ± 0.1§ | 27 ± 5 § (30) |
| **4F** | 1.5 ± 0.2§ | 11 ± 2§ (7.5) |
| **5F** | 0.6 ± 0.3§ | 26 ± 1§ (42) |
| **6F** | 1.3 ± 0.2§ | 8 ± 2§ (6.0) |
| **7F**§ | 2.3 ± 0.2§ | 16 ± 1 (7.0) |

§Co-crystal structures determined in this work. †Co-crystal structures previously determined by our laboratory (Soumana D.I. et al, *ACS Chem Biol*, 11(4):900, 2016, Romano K.P. et al, *Plos Pathog*, 8(7):e1002832, 2012, and (Matthew N.M. et al, *J. Med. Chem*. 60(13):5699, 2017). (Matthew N.M., Zephyr J. et al, *mBio* 60(13):5699, 2020)

**Table S2.** X-ray data collection and crystallographic refinement statistics.

| **Complex structures** | **WT1a 3** | **WT1a 4** | **WT1a 5** | **WT1a 1F** | **WT1a 2F*S*** | **WT1a 2F*R*** | **WT1a 3F** | **WT1a 5F** | **WT1a 6F** | **WT1a 7F** |
|---|---|---|---|---|---|---|---|---|---|---|
| **PDB ID** | 7MM4 | 7MM2 | 7MM3 | 6DIS | 7MM7 | 7MMA | 7MM6 | 7MM9 | 7MM5 | 7MM8 |
| **Resolution (Å)** | 1.89 | 1.89 | 1.78 | 1.92 | 1.86 | 1.65 | 2.00 | 2.11 | 1.90 | 1.43 |
| **Space group** | $P2_12_12_1$ | $P2_12_12_1$ | $P2_12_12_1$ | $P2_12_12_1$ | $P2_12_12_1$ | $P2_12_12_1$ | $P2_12_12_1$ | $P2_12_12_1$ | $P2_12_12_1$ | $P2_12_12_1$ |
| **Molecules in AU**[a] | 1 | 1 | 1 | 1 | 1 | 1 | 1 | 1 | 1 | 1 |
| **Cell dimensions:** | | | | | | | | | | |
| **a (Å)** | 55.0 | 54.5 | 54.9 | 55.6 | 54.7 | 54.1 | 54.6 | 54.6 | 54.1 | 45.4 |
| **b (Å)** | 58.7 | 58.7 | 58.7 | 58.6 | 58.7 | 58.5 | 58.7 | 58.6 | 58.4 | 59.2 |
| **c (Å)** | 59.8 | 59.5 | 59.9 | 60.0 | 59.7 | 59.7 | 59.7 | 60.0 | 61.4 | 96.7 |
| **β (°)** | 90 | 90 | 90 | 90 | 90 | 90 | 90 | 90 | 90 | 90 |
| **Completeness (%)** | 99.5 | 99.8 | 99.1 | 96.7 | 99.7 | 93.2 | 99.1 | 99.6 | 96.5 | 97.7 |
| **Total reflections** | 110340 | 89940 | 116617 | 106822 | 112740 | 115326 | 88404 | 77244 | 108482 | 175070 |
| **Unique reflections** | 15968 | 15783 | 18600 | 14999 | 16565 | 21869 | 13328 | 11516 | 15462 | 47936 |
| **Average I/σ** | 13.4 | 12.9 | 18.6 | 19.8 | 15.5 | 25.3 | 12.9 | 23.8 | 17.0 | 16.6 |
| **Redundancy** | 6.9 | 5.7 | 6.3 | 7.1 | 6.8 | 5.3 | 6.6 | 6.7 | 7.0 | 3.7 |
| $R_{sym}$ **(%)**[b] | 12.9 (38.4) | 10.0 (41.3) | 6.68 (25.6) | 10.6 (45.8) | 9.2 (29.8) | 4.5 (12.5) | 11.2 (37.0) | 8.0 (24.4) | 7.9 (35.8) | 5.7 (32.1) |
| **RMSD**[c] **in:** | | | | | | | | | | |
| **Bond lengths (Å)** | 0.006 | 0.005 | 0.011 | 0.01 | 0.009 | 0.011 | 0.003 | 0.006 | 0.006 | 0.021 |
| **Bond angles (°)** | 0.92 | 0.75 | 1.14 | 1.0 | 1.04 | 1.19 | 0.54 | 0.84 | 0.84 | 1.82 |
| $R_{factor}$ **(%)**[d] | 16.1 | 16.0 | 15.1 | 18.7 | 15.4 | 14.4 | 17.5 | 17.7 | 20.0 | 18.5 |
| $R_{free}$ **(%)**[e] | 20.4 | 20.0 | 18.9 | 22.7 | 18.1 | 18.1 | 19.7 | 21.0 | 23.9 | 20.4 |

**Table S2 (continued).** X-ray data collection and crystallographic refinement statistics.

| Complex structures | D168A 1 | D168A 3 | D168A 5 | D168A 7 | D168A 1F | D168A 2$F_S$ | D168A 2$F_R$ | D168A 3F | D168A 4F | D168A 5F | D168A 6F | D168A 7F |
|---|---|---|---|---|---|---|---|---|---|---|---|---|
| **PDB ID** | 7MM6 | 7MMC | 7MMB | 7MMD | 6DIW | 7MMI | 7MML | 7MMH | 7MMG | 7MMK | 7MMF | 7MMJ |
| **Resolution (Å)** | 1.56 | 2.00 | 1.99 | 1.89 | 1.80 | 1.80 | 1.70 | 1.75 | 1.95 | 1.89 | 1.89 | 1.89 |
| **Space group** | $P2_12_12_1$ | $P2_12_12_1$ | $P2_12_12_1$ | $P2_12_12_1$ | $P2_12_12_1$ | $P2_12_12_1$ | $P2_12_12_1$ | $P2_12_12_1$ | $P12_11$ | $P2_12_12_1$ | $P2_12_12_1$ | $P2_12_12_1$ |
| **Molecules in AU**[a] | 1 | 1 | 1 | 1 | 1 | 1 | 1 | 1 | 2 | 1 | 1 | 1 |
| **Cell dimensions:** | | | | | | | | | | | | |
| **a (Å)** | 55.6 | 45.3 | 54.3 | 55.1 | 55.7 | 54.0 | 54.0 | 54.9 | 39.9 | 54.4 | 54.8 | 55.2 |
| **b (Å)** | 58.5 | 58.9 | 58.7 | 58.7 | 58.6 | 58.6 | 58.6 | 58.7 | 51.0 | 58.6 | 59.8 | 58.7 |
| **c (Å)** | 59.9 | 96.2 | 59.5 | 59.9 | 60.1 | 59.6 | 59.6 | 58.8 | 77.1 | 59.6 | 58.7 | 59.8 |
| **β (°)** | 90 | 90 | 90 | 90 | 90 | 90 | 90 | 90 | 90,93,90 | 90 | 90 | 90 |
| **Completeness (%)** | 98.3 | 96 | 99.8 | 85.9 | 92.6 | 94.4 | 99.8 | 99.8 | 100 | 99.6 | 99.8 | 85.9 |
| **Total reflections** | 249313 | 104573 | 91993 | 55093 | 120162 | 94587 | 109184 | 119179 | 144992 | 108115 | 109121 | 55093 |
| **Unique reflections** | 28103 | 17308 | 13596 | 13836 | 17476 | 17155 | 21349 | 20078 | 22738 | 15843 | 15907 | 13836 |
| **Average I/σ** | 25.2 | 15.3 | 21.3 | 12.1 | 6.3 | 15.1 | 19.7 | 19.3 | 7.6 | 27.5 | 17.0 | 12.1 |
| **Redundancy** | 8.9 | 6.0 | 6.8 | 4.0 | 6.9 | 5.5 | 5.1 | 5.9 | 6.4 | 6.8 | 6.9 | 4.0 |
| $R_{sym}$ **(%)**[b] | 4.7 (14.3) | 7.3 (23.5) | 8.8 (29.9) | 7.5 (26.4) | 3.9 (11.7) | 7.6 (43.4) | 5.9 (31.1) | 6.2 (32.6) | 16.7 (41.8) | 6.9 (28.9) | 7.9 (35.8) | 7.5 (26.4) |
| **RMSD**[c] **in:** | | | | | | | | | | | | |
| **Bond lengths (Å)** | 0.015 | 0.005 | 0.007 | 0.010 | 0.010 | 0.006 | 0.015 | 0.008 | 0.010 | 0.005 | 0.012 | 0.014 |
| **Bond angles (°)** | 1.6 | 0.63 | 1.05 | 0.83 | 1.5 | 0.83 | 1.4 | 1.04 | 1.03 | 1.12 | 1.07 | 1.52 |
| $R_{factor}$ **(%)**[d] | 14.4 | 18.6 | 18.3 | 15.7 | 13.8 | 17.2 | 15.9 | 15.7 | 19.8 | 16.2 | 16.5 | 19.5 |
| $R_{free}$ **(%)**[e] | 17.0 | 21.0 | 22.3 | 19.9 | 18.0 | 20.8 | 19.3 | 19.9 | 24.9 | 20.8 | 19.9 | 23.9 |

[a]AU, asymmetric unit.

[b]$R_{sym} = \Sigma \mid I - \langle I \rangle \mid / \Sigma I$, where $I$ = observed intensity, $\langle I \rangle$ = average intensity over symmetry equivalent; values in parentheses are for the highest resolution shell.

[c]RMSD, root mean square deviation.

[d]$R_{factor} = \Sigma \| F_o| - |F_c\| / \Sigma |F_o|$.

[e]$R_{free}$ was calculated from 5% of reflections, chosen randomly, which were omitted from the refinement process
